# Subtype-specific secretion of extracellular vesicles by LRRK2 and Rab GTPases under lysosomal stress

**DOI:** 10.64898/2026.04.14.718125

**Authors:** Maria Sakurai, Tomoki Kuwahara, Shoichi Suenaga, Sho Takatori, Taisuke Tomita, Tammy Shalit, Elizabeth Tengstrand, Frank Hsieh, Takeshi Iwatsubo

**Affiliations:** Department of Neuropathology, Graduate School of Medicine, The University of Tokyo, 7-3-1 Hongo, Bunkyo-ku, Tokyo, 113-0033, Japan; Laboratory of Neuropathology and Neuroscience, Graduate School of Pharmaceutical Sciences, The University of Tokyo, 7-3-1 Hongo, Bunkyo-ku, Tokyo, 113-0033, Japan; Nextcea Inc, 500 West Cummings Park, Suite 4550, Woburn, MA, 01801, United States; Department of Dementia Inclusion and Therapeutics, The University of Tokyo Hospital, 7-3-1 Hongo, Bunkyo-ku, Tokyo, 113-0033, Japan; National Center of Neurology and Psychiatry, 4-1-1 Ogawa-Higashi, Kodaira, Tokyo, 187-8551, Japan

**Keywords:** Extracellular vesicles, LRRK2, Rab, lysosomal stress, Parkinson’s disease

## Abstract

LRRK2, the Parkinson’s disease-associated kinase, phosphorylates a subset of Rab GTPases and regulates membrane dynamics. We previously reported that lysosomal stress activates LRRK2 and thereby induces the exocytic secretion of lysosomal contents, but the detailed secretion mechanism remained unclear. Here we found that, under lysosomal stress, endolysosomal luminal and membrane components were secreted with extracellular vesicles (EVs) via LRRK2. Bis(monoacylglycerol)phosphate, an endolysosomal lipid and a urinary marker of LRRK2 activity, was similarly secreted via LRRK2, whereas CD9-positive EVs were not involved. Further dissection of the secreted EVs revealed that Alix-positive EVs were secreted via Rab8a as well as the ESCRT component VPS4, whereas LAMP1/cathepsin B-positive EVs were secreted via Rab10/Rab35, and SNARE proteins syntaxin 2 and VAMP8 regulated the secretion of both EV subtypes. These findings suggest a distinctive stress-induced secretory mechanism whereby LRRK2 facilitates the secretion of multiple EV subtypes by controlling Rab GTPases involved in each pathway.

## Introduction

Leucine rich repeat kinase 2 (*LRRK2*) is the well-known causative gene for autosomal dominant Parkinson’s disease (PD)^1–3^ as well as one of top risk genes for sporadic PD^4–6^. *LRRK2* variants are also associated with the risk of several inflammatory or infectious diseases, such as Crohn’s disease^7,8^ and leprosy^9,10^. LRRK2 is highly expressed in various tissues and cells including brain, kidney, lung and immune cells^11^. Among these, the role of LRRK2 in macrophages and microglia is of particular interest as the expression of LRRK2 in these cells is potently induced by inflammatory stimuli^12–15^. Also, PD-associated genetic variants have been shown to influence LRRK2 expression in microglia^16^. *LRRK2* encodes a serine/threonine kinase that phosphorylates a subset of Rab GTPases, such as Rab8a, Rab10 and Rab35, in cells^17^. Since Rab GTPases generally regulate intracellular membrane trafficking and their phosphorylation sites by LRRK2 lie in the middle of the switch II region critical for their function (*e.g.*, Thr73 in Rab10), LRRK2 has been postulated to modify Rab-related membrane trafficking pathways. Importantly, PD-associated mutations in LRRK2 commonly enhance its activity to phosphorylate Rab GTPases^17,18^, suggesting the possible involvement of Rab dysregulation in the pathogenesis of PD.

Recent studies on LRRK2 in membrane trafficking have focused particularly on lysosomes. LRRK2 deficient animals exhibit an age-dependent accumulation of enlarged secondary lysosomes or lysosome-related organelles in the kidney and lung^19–21^. Di-22:6-bis(monoacylglycerol)phosphate (BMP), a negatively charged phospholipid enriched in late endosomal and lysosomal membranes^22,23^, has also been shown to accumulate in the kidney parenchyma of LRRK2 deficient mice^24^, while its level in urine is significantly decreased^21,24^. The decrease in urinary BMP levels is also clearly detected in humans treated with LRRK2 inhibitors in clinical trials, establishing it as an indicator for assessing the *in vivo* efficacy of LRRK2 inhibitors^25^. Urinary BMP is now considered to be associated with PD, as its levels have been shown to be elevated in both LRRK2-asocciated and idiopathic PD^26–28^.

In studies using cultured cells, we have previously reported that treatment of RAW264.7 macrophage cells with the lysosomotropic agent chloroquine (CQ) causes LRRK2 activation and its recruitment onto the enlarged lysosomes^29^. Subsequent studies have reported that other lysosomal membrane-targeted drugs, such as L-leucyl-L-leucine methyl ester (LLOMe) and monensin, also cause LRRK2 activation and lysosomal recruitment^30–32^. We additionally reported that CQ-induced lysosomal stress induces exocytic release of lysosomal enzymes such as mature cathepsin B/D via LRRK2 and Rab10 in RAW264.7 cells^29^. This LRRK2/Rab10-mediated release was also involved in the release of insoluble α-synuclein from microglia loaded with α-synuclein pre-formed fibrils ^33^. Furthermore, a recently established mechanism termed conjugation of ATG8 to single membranes (CASM), which involves non-autophagic role of the ATG8 conjugation system consisting mainly of the ATG12–ATG5-ATG16L1 complex, was responsible for the activation of LRRK2 and the resultant release of lysosomal contents^34^. Thus, LRRK2, Rab10 and the ATG8 conjugation system have been identified as mediators of exocytic release under lysosomal stress, although more detailed mechanism of the release remained unclear.

Extracellular vesicles (EVs) are cell-derived small vesicular membranes, including exosomes and microvesicles. A recent study has reported that the secretion of exosome-like EVs as well as secretory autophagy are upregulated in neurons expressing G2019S pathogenic mutant LRRK2^35^. Additionally, LRRK2 has been implicated in the secretion of EVs containing BMP^24,36^, while direct evidence of LRRK2 regulating the secretion of BMP vesicles has not been provided. EVs are topologically assumed to enclose cytoplasmic molecules. In contrast, LRRK2-mediated secretion we observed under lysosomal stress, as described above, primarily involves the secretion of lysosomal luminal enzymes and is evident in macrophage lineage cells such as RAW264.7 cells and microglial cells. Therefore, given the role of LRRK2 in PD, it was desired to further characterize this LRRK2-mediated secretion in comparison with known secretory mechanisms.

In this study, we found that lysosomal hydrolases secreted by LRRK2 under stress were indeed present in the EV fraction, attached on their surface. Subsequent analyses revealed that LRRK2 facilitates the secretion of a subset of EVs with endolysosomal components, including BMP, and that each LRRK2-substrate Rab regulates the secretion of distinct EV subtypes. Thus, the broader role of LRRK2 in EV secretion under lysosomal stress has become apparent.

## Results

### Lysosomal stress induces LRRK2-mediated secretion of endolysosomal components into EV fractions

To identify the secretory pathway regulated by LRRK2, we first examined the possible involvement of three known pathways that differ in membrane dynamics: lysosomal exocytosis, secretory autophagy, and EV secretion **(Fig. S1A)**^37,38^. All these pathways have the potential to secrete lysosomal components and involve ATG8 conjugation system proteins that act upstream of LRRK2-mediated release^29^. We used RAW264.7 mouse macrophage cells for analysis, as this cell line exhibits high LRRK2 expression and secretory activity under lysosomal stress. In lysosomal exocytosis, leakage of calcium ions from lysosomes triggers their fusion with plasma membrane, releasing their contents^39–41^. We confirmed that the treatment with ionomycin, a calcium ionophore known to induce lysosomal exocytosis, increased the extracellular activity of the lysosomal β-hexosaminidase (β-hex), but this effect was not suppressed by knockdown of LRRK2, ruling out the involvement of LRRK2 in the lysosomal exocytosis **(Fig. S1B)**. Secretory autophagy, the second pathway, employs double-membrane autophagosomes to release their contents^42^. Although very few markers are known that discriminate it from other pathways, interleukin-1β (IL-1β) was suggested to be released through secretory autophagy upon treatment with nigericin, an NLRP3 inflammasome activator^43–45^. We confirmed that nigericin treatment increased the extracellular level of IL-1β, but this was not suppressed by treatment with the LRRK2 inhibitor MLi-2 **(Fig. S1C)**, ruling out the involvement of LRRK2 in secretory autophagy.

The secretion of EVs, the third pathway, involves diverse cell-derived vesicles. Those originating from endolysosomes are primarily exosomes generated by their invagination to form intraluminal membranes, with Alix being a representative marker. Analysis of EV fractions recovered by ultracentrifugation of the culture medium revealed that, upon exposure to CQ, the levels of not only Alix but also the lysosomal membrane protein LAMP1 and the lysosomal luminal enzymes cathepsin B/D (mature and intermediate mature forms) were increased in EV fractions **(Fig. 1A-D)**. Cathepsin B/D were also collected in the supernatant fractions after ultracentrifugation **(Fig. 1A and 1D)**. Importantly, CQ-induced secretion of these proteins into EV fractions was suppressed by LRRK2 inhibitor treatment **(Fig. 1E-H)** and indeed dependent on LRRK2 kinase activity **(Fig. S2A)**. The activity of lysosomal β-hex in EV fractions as well as media was also increased upon CQ treatment, and this increase was again suppressed by LRRK2 kinase inhibition **(Fig. 1I and 1J)**. The absence of passive leakage due to cell death or membrane damage upon CQ treatment was confirmed by the lack of increased lactate dehydrogenase (LDH) activity in the culture media **(Fig. S3A)**. These results indicate that lysosomal luminal and membrane proteins as well as Alix are all secreted with EVs under the control of LRRK2.

**Figure 1.**
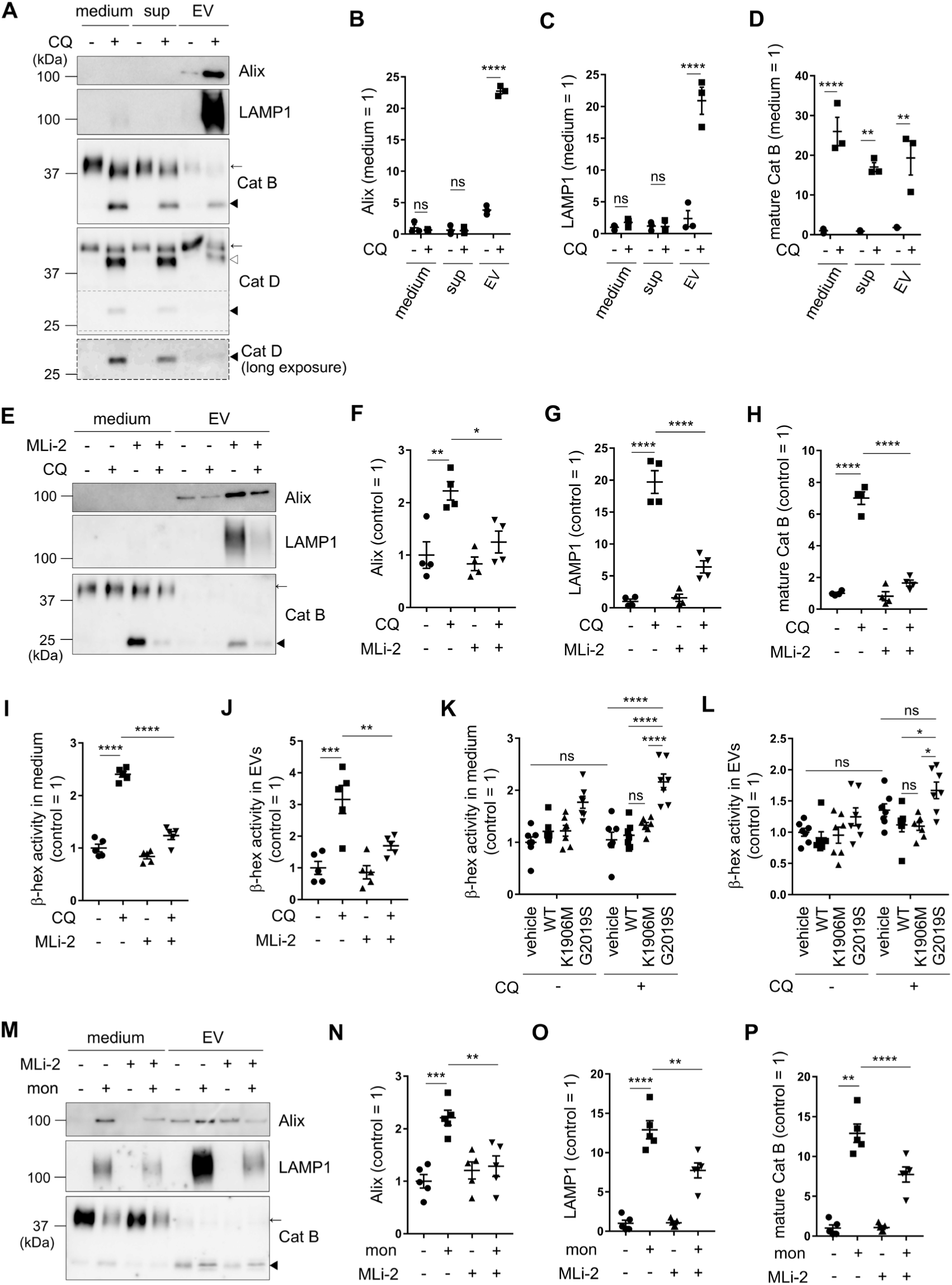
Lysosomal stress induces LRRK2-mediated secretion of endolysosomal components into EV fractions. **(A)** Immunoblotting of medium, sup and EV fractions derived from RAW264.7 cells treated with or without chloroquine (CQ). medium: fractions before ultracentrifugation, sup: supernatants after ultracentrifugation, EV: pellets after ultracentrifugation. Closed filled arrowheads indicate mature cathepsin B (Cat B) and cathepsin D (Cat D), a closed open arrowhead indicates intermediate mature Cat D, and arrows indicate immature Cat B and Cat D. **(B-D)** Quantification of Alix (B), LAMP1 (C) and mature cathepsin B (D) in each fraction, as shown in A. n = 3, mean ± SEM, two-way ANOVA with Tukey’s test. ** *p* < 0.01, **** *p* < 0.0001. **(E)** Immunoblotting of medium and EV fractions derived from cells treated with or without MLi-2 and CQ. **(F-H)** Quantification of Alix (F), LAMP1 (G) and mature Cat B (H) in EV fractions, as shown in E. n = 4, mean ± SEM, one-way ANOVA with Tukey’s test. * *p* < 0.05, ** *p* < 0.01, **** *p* < 0.0001. **(I, J)** β-Hexosaminidase (β-hex) activity in medium (I) and EV (J) fractions derived from cells treated with or without CQ and MLi-2. n = 5, mean ± SEM, one-way ANOVA with Tukey’s test. ** *p* < 0.01, *** *p* < 0.001, **** *p* < 0.0001. **(K, L)** β-Hex activity in medium (K) and EV (L) fractions derived from CQ-treated or untreated HEK293 cells stably overexpressing the indicated mutant LRRK2. n = 7, mean ± SEM, two-way ANOVA with Tukey’s test. * *p* < 0.05, **** *p* < 0.0001. **(M)** Immunoblotting of medium and EV fractions derived from cells treated with or without monensin (mon) and MLi-2. **(N-P)** Quantification of Alix (N), LAMP1 (O) and mature Cat B (P) in EV fractions, as shown in M. n = 4, mean ± SEM, one-way ANOVA with Tukey’s test. ** *p* < 0.01, *** *p* < 0.001, **** *p* < 0.0001.

Since β-hex activity assay enables detection of even small increases in secretion, we investigated whether CQ-dependent secretion of EVs occurs in cells with lower secretory activity compared to macrophage lineage cells. Analysis of HEK293 cells revealed that the secreted β-hex in media upon CQ treatment was higher in cells overexpressing PD-associated mutant (G2019S) LRRK2 compared to cells overexpressing wild-type or kinase-dead (K1906M) LRRK2, as well as cells that did not overexpress LRRK2 **(Fig. 1K)**. β-Hex activity in EV fractions was also higher in HEK293 cells overexpressing G2019S LRRK2 compared to cells overexpressing wild-type or K1906M LRRK2 **(Fig. 1L)**. This may suggest that LRRK2-dependent enhanced EV secretion could also be involved in the pathogenetic process of PD.

To further analyze the effects of lysosomal stress stimuli other than CQ, we tested the effect of monensin, an ionophore that targets lysosomal membranes and also activates LRRK2^32^. Monensin treatment induced the secretion of Alix, LAMP1 and mature cathepsin B into EV fractions in a LRRK2 kinase activity-dependent manner **(Fig. 1M-P, Fig. S2A)**, confirming the role of LRRK2 in broader lysosomal stress. Since CQ exhibited a stronger effect on EV secretion than monensin, CQ was used in subsequent analyses.

### LRRK2 regulates the secretion of exosome-like EVs attaching lysosomal enzymes

Next, to confirm that the secreted EVs were generated via the exosome biosynthesis pathway, we treated cells with or without GW4869 under CQ stimulation. GW4869 is a broad-spectrum inhibitor of exosome biogenesis that primarily targets neutral sphingomyelinase 2 (nSMase2) but also effects on Alix-containing exosomes^46^. CQ-induced secretion of Alix and mature cathepsin B was suppressed by GW4869 **(Fig. 2A-D)**, supporting the secretion of exosome-like vesicles. Rab10 phosphorylation was not suppressed by GW4869 treatment, indicating that GW4869 inhibits EV secretion independently of LRRK2 **(Fig. S2B)**. We then performed direct observation of the secreted vesicles in the culture media using electron microscopy and found that vesicles with a diameter of ∼100 nm, consistent with exosome-like small EVs, were abundant in the CQ-treated samples **(Fig. 2E)**. We further investigated changes in the number and size of secreted EV particles upon CQ and/or LRRK2 inhibitor treatment (MLi-2) by performing nanoparticle tracking analysis (NTA). NTA revealed that the particle size was comparable in all samples at <150 nm **(Fig. 2F and 2H)**, while the number of particles increased significantly upon CQ treatment, and this increase was suppressed by LRRK2 inhibition **(Fig. 2F and 2G)**. These results were consistent with the biochemical analyses conducted thus far, suggesting the role of LRRK2 in the secretion of exosome-like EVs.

**Figure 2.**
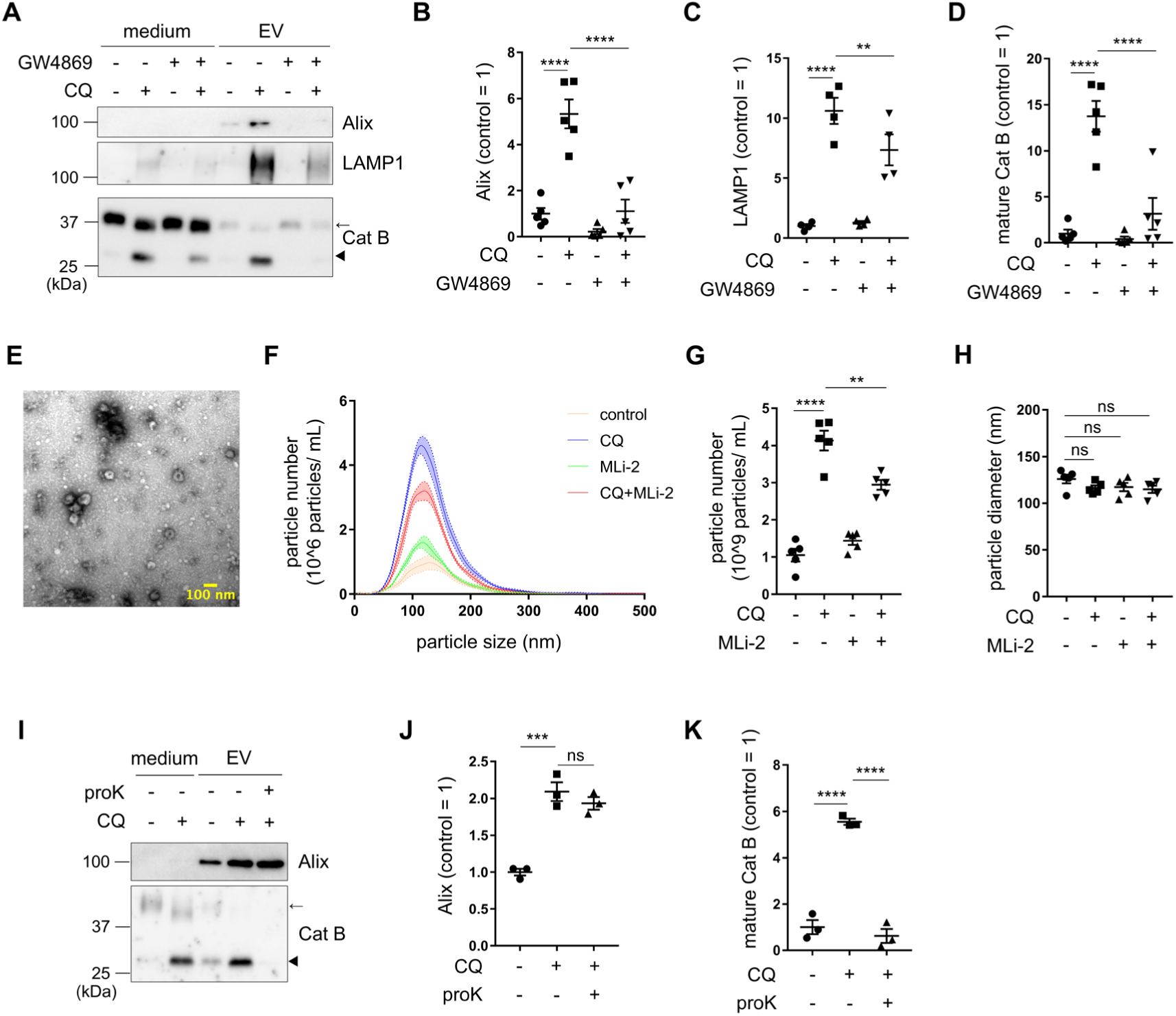
LRRK2 regulates the secretion of exosome-like EVs attaching lysosomal enzymes. **(A)** Immunoblotting of medium and EV fractions derived from RAW264.7 cells treated with or without GW4869 and CQ. Arrowhead: mature cathepsin B (Cat B), arrow: immature Cat B. **(B-D)** Quantification of Alix (B), LAMP1 (C) and mature Cat B (D) in EV fractions. n = 5, mean ± SEM, one-way ANOVA with Tukey’s test. ** *p* < 0.01, **** *p* < 0.0001. **(E)** Electron micrographs of EVs secreted in culture medium. Scale bar: 100 nm. **(F)** Size and number of the secreted EV particles measured by NTA. **(G, H)** Quantification of the particle number (G) and mode of particle size (H)detected in the culture media from cells treated with or without CQ and MLi-2, as shown in F. n = 5, mean ± SEM, one-way ANOVA with Tukey’s test. ** *p* < 0.01, **** *p* < 0.0001. **(I)** Immunoblotting of medium and EV fractions derived after proteinase K (proK) treatment of the medium from CQ-treated cells. Arrowhead: mature Cat B, arrow: immature Cat B. **(J, K)** Quantification of Alix (J) and mature Cat B (K) in EV fractions, as shown in I. n = 4, mean ± SEM, one-way ANOVA with Tukey’s test. *** *p* < 0.001, **** *p* < 0.0001.

Since exosomes are generated by invagination of the endolysosomal membranes and topologically enclose cytoplasmic components, how lysosomal luminal proteins such as cathepsin B are secreted with exosome-like EVs remained unclear. On the other hand, recent studies have raised the possibility that lysosomal enzymes are released by attaching to the outer surface of EVs^47,48^. To determine the localization of lysosomal enzymes relative to EVs, we performed digestion assays by adding proteinase K to the culture medium containing EVs. As expected, mature cathepsin B was digested by proteinase K, while Alix, a cytoplasmic protein present inside EVs, remained intact **(Fig. 2I-K)**. These results together suggest that EVs attaching lysosomal enzymes are secreted via LRRK2 under lysosomal stress.

### LRRK2 regulates the secretion process rather than the formation of EVs

To clarify the site of action of LRRK2 in the process leading to the secretion of exosome-like EVs, we next examined the possibility that LRRK2 regulates the process of vesicle formation. Intraluminal vesicles (ILVs) in RAW264.7 cells treated with CQ were observed using electron microscopy, revealing the presence of ILVs of various sizes within enlarged lysosomes **(Fig. 3A)**. In cells treated with both CQ and the LRRK2 inhibitor MLi-2, we observed more enlarged lysosomes, consistent with our previous reports^29,34^ **(Fig. 3A-C)**, and the number of ILVs within these lysosomes was rather increased compared to cells treated with CQ only **(Fig. 3D)**. The ratio of ILV count in lysosomes to lysosomal area showed no significant differences between MLi-2-treated cells and untreated cells **(Fig. 3E)**. These data suggest that LRRK2 does not affect vesicle formation of EVs but instead regulates the secretion process after formation.

**Figure 3.**
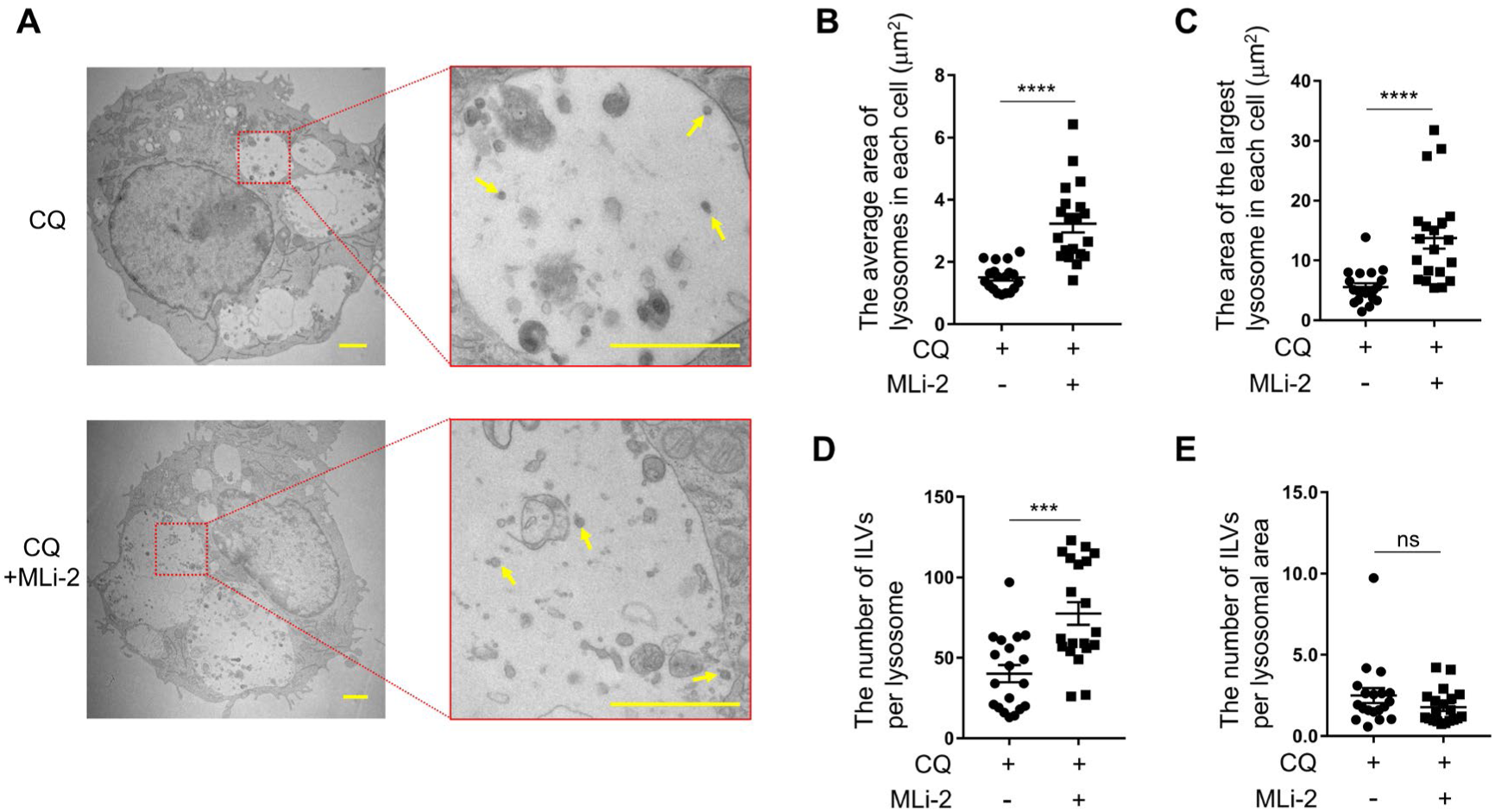
LRRK2 regulates the secretion process rather than the formation of EVs. **(A)** Electron micrographs of RAW264.7 cells treated with CQ or CQ plus MLi-2. Scale bar: 1 μm. Arrows indicate representative ILVs in each lysosome. **(B)** Quantification of the average area of lysosomes in each cell. n = 20, mean ± SEM, unpaired *t*-test. **** *p* < 0.0001. **(C)** Quantification of the area of the largest lysosome in each cell. n = 20, mean ± SEM, unpaired *t*-test. **** *p* < 0.0001. **(D)** Quantification of the number of ILVs per lysosome. n = 20, mean ± SEM, unpaired *t*-test. *** *p* < 0.001. **(E)** Quantification of the number of ILVs per lysosomal area. n = 20, mean ± SEM, unpaired *t*-test.

### EV membranes secreted by LRRK2 contain di-22:6-BMP but not CD9

Di-22:6-BMP is a unique phospholipid specifically enriched in endolysosomal intralumenal membranes, and its level in urine is well correlated with LRRK2 kinase activity *in vivo*. Since LRRK2-regulated EVs contained lysosomal components, we hypothesized that di-22:6-BMP is also contained in EVs secreted via LRRK2 under lysosomal stress. We utilized mass spectrometry to measure the amount of di-22:6 BMP in EV fractions as well as cell pellets from RAW264.7 cells treated with CQ and/or MLi-2. The BMP level in EV fraction was markedly increased upon CQ treatment, and this increase was significantly suppressed by additional MLi-2 treatment **(Fig. 4A)**. BMP in cell pellets was decreased upon CQ treatment regardless of MLi-2 treatment **(Fig. 4B)**, possibly due to impaired endolysosomal maturation under CQ-treated conditions^49^. These data suggest that LRRK2 is involved in the secretion of BMP-containing EVs under lysosomal stress.

**Figure 4.**
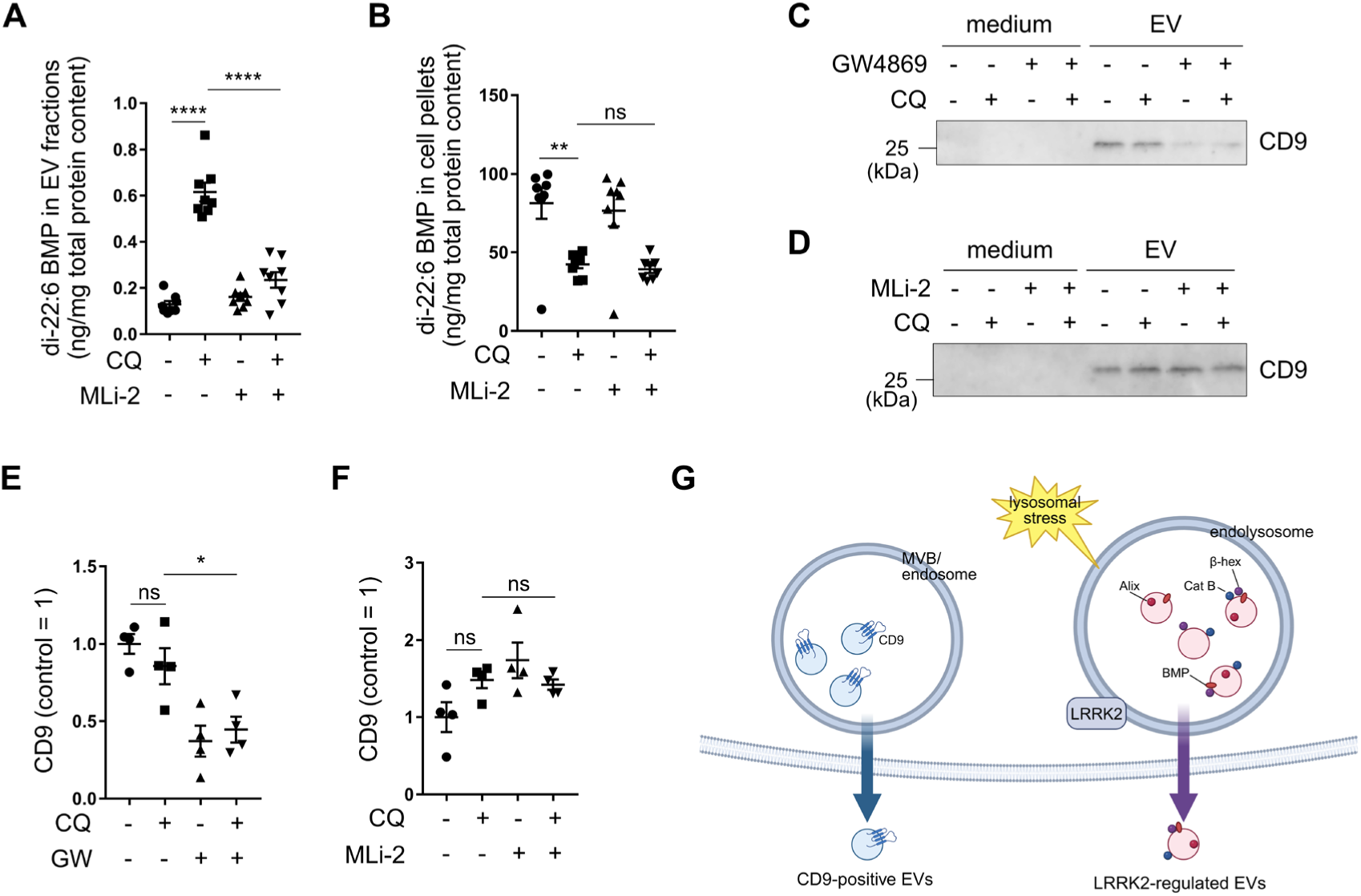
EV membranes secreted by LRRK2 contain di-22:6-BMP but not CD9. **(A)** The amount of di-22:6-BMP normalized by total protein content in EV fractions derived from RAW264.7 cells treated with or without CQ and MLi-2. n = 4, mean ± SEM, one-way ANOVA with Tukey’s test. ** *p* < 0.01. **(B)** The amount of di-22:6-BMP normalized by total protein content in cell pellets derived from cells treated with or without CQ and MLi-2. n = 4, mean ± SEM, one-way ANOVA with Tukey’s test. **** *p* < 0.0001. **(C, D)** Immunoblot analysis of CD9 in medium and EV fractions derived from RAW264.7 cells treated with or without CQ plus GW4869 (C) or MLi-2 (D). **(E, F)** Quantification of CD9 in EV fractions from cells treated with or without CQ plus GW4869 (GW) (E) or MLi-2 (F), as shown in C and D. n = 4, mean ± SEM, one-way ANOVA with Tukey’s test. * *p* < 0.05. **(G)** Model of LRRK2-dependent and LRRK2-independent secretion of EVs with different membrane compositions.

We further characterized the components of EV membranes secreted by LRRK2. The tetraspanin family proteins such as CD9 and CD63 are enriched on a subset of exosome-like EV membranes and thus have been regarded as representative EV membrane markers. Since expression of CD63 was hardly detected in our RAW264.7 cells, we analyzed the secretion of CD9 into the EV fraction. We found that, unlike other membrane components such as BMP and LAMP1, no increase of CD9 was observed upon CQ treatment **(Fig. 4C-F)**. Also, CD9 in EVs was decreased by GW4869 treatment but not by MLi-2 treatment **(Fig. 4C-F)**. These results suggest that LRRK2 facilitates the secretion of a subset of EVs that do not contain CD9-positive EVs **(Fig. 4G)**.

### VPS4 and Rab8a regulate the secretion of Alix-positive EVs under lysosomal stress

Among exosome-like EVs, tetraspanin-negative EVs are thought to be mostly generated by the endosomal sorting complexes required for transport (ESCRT) machinery, which facilitates ILV formation by inducing endolysosomal membrane invagination. To analyze the involvement of ESCRT in LRRK2-mediated EV secretion, we performed knockdown of *Vps4*, an ESCRT-Ⅲ interactor essential for the final step of membrane invagination. Unexpectedly, under CQ-treated conditions, *Vps4* knockdown caused a decrease in the amount of Alix in secreted EVs, while LAMP1 and mature cathepsin B were not decreased but rather increased, potentially reflecting a compensatory effect **(Fig. 5A-D, Fig. S4A)**. The absence of cytotoxicity upon knockdown was additionally confirmed by LDH activity assays **(Fig. S3B)**. This suggests that Alix-positive EVs and LAMP1/cathepsin B-positive EVs are different and are regulated separately.

**Figure 5.**
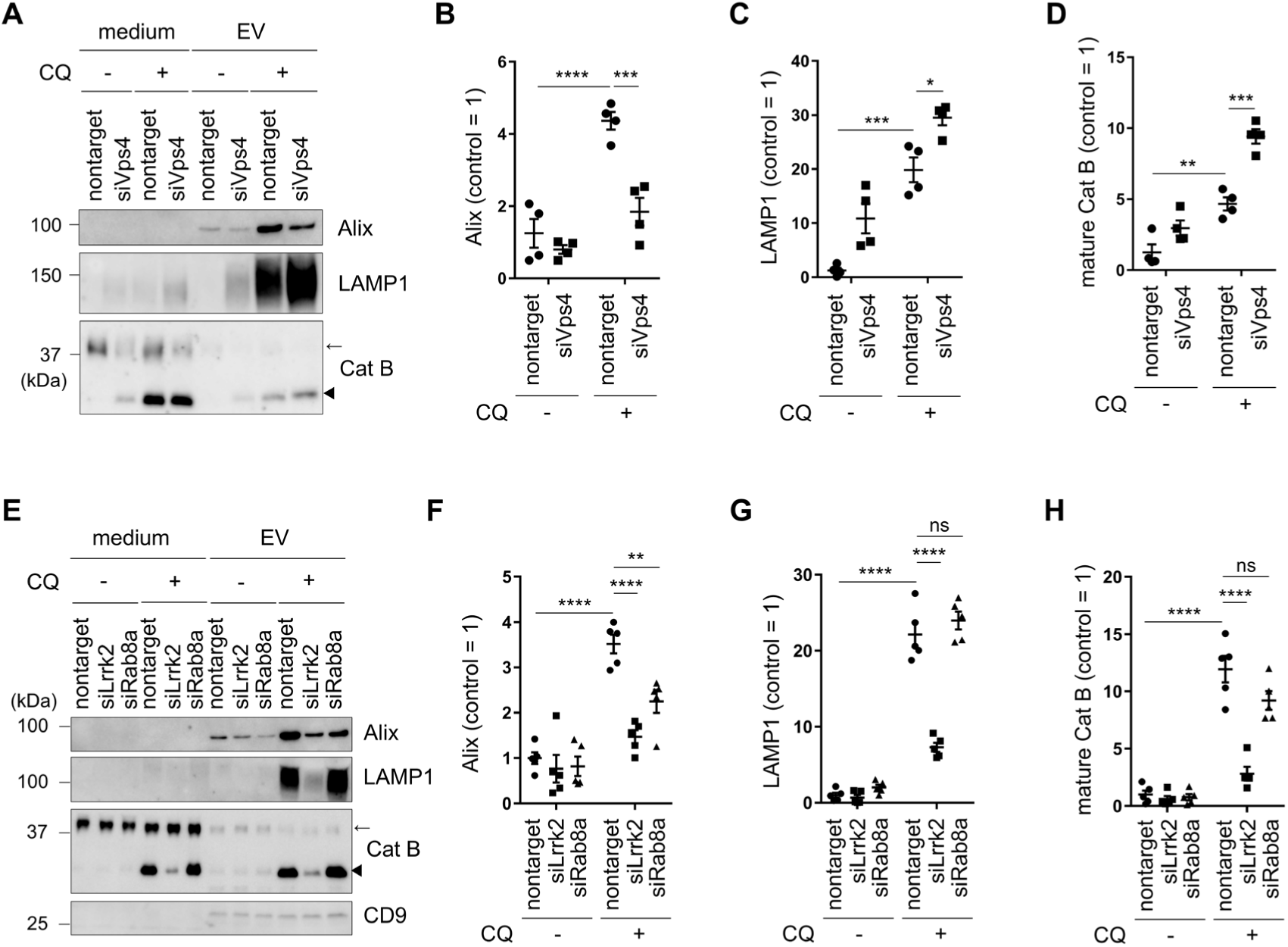
VPS4 and Rab8a regulate the secretion of Alix-positive EVs under lysosomal stress. **(A)** Immunoblotting of medium and EV fractions derived from RAW264.7 cells treated with nontarget or VPS4 siRNAs. Arrowhead: mature cathepsin B (Cat B), arrow: immature Cat B. **(B-D)** Quantification of Alix (B), LAMP1 (C) and mature Cat B (D) in EV fractions. n = 4, mean ± SEM, two-way ANOVA with Tukey’s test. * *p* < 0.05, ** *p* < 0.01, *** *p* < 0.001, **** *p* < 0.0001. **(E)** Immunoblotting of medium and EV fractions derived from cells treated with LRRK2 or Rab8a siRNAs. Arrowhead: mature Cat B, arrow: immature Cat B. **(F-H)** Quantification of Alix (F), LAMP1 (G) and mature Cat B (H) in EV fractions. n = 5, mean ± SEM, two-way ANOVA with Tukey’s test. ** *p* < 0.01, *** *p* < 0.001, **** *p* < 0.0001.

We also focused on the involvement of Rab8a, one of LRRK2 substrates, because Rab8a has been implicated in the secretion of exosomes^50,51^. *Rab8a* knockdown under CQ-treated conditions caused a decrease in Alix levels in EVs while LAMP1 and mature cathepsin B were unchanged, unlike the effects on *Lrrk2* knockdown **(Fig. 5E-H, Fig. S4B)**. Collectively, these data suggest that Rab8a and the ESCRT machinery are responsible for the secretion of Alix-positive EVs that are thought to originate from late endosomal compartments but not LAMP1/cathepsin B-positive EVs originating from lysosomes.

### Rab10 and Rab35 regulate the secretion of LAMP1/cathepsin B-positive EVs under lysosomal stress

We also investigated the role of LRRK2 substrate Rab GTPases other than Rab8a in EV secretion. We have previously shown that Rab10 is involved in the release of cathepsins into the medium upon CQ treatment^29,32,34^. Here we focused on Rab10 as well as Rab35, the latter being implicated in the secretion of exosomes^52^. Knockdown of *Rab10* or *Rab35* caused a decrease in the secretion of LAMP1/cathepsin B-positive EVs upon CQ treatment, whereas the secretion of Alix-positive EVs was unaffected by *Rab35* knockdown and rather increased by *Rab10* knockdown **(Fig. 6A-D, Fig. S4C)**. The absence of cytotoxicity upon knockdown was again confirmed **(Fig. S3B)**. This result suggests that Rab10 and Rab35 regulate the secretion of only LAMP1/cathepsin B-positive EVs of lysosomal origin, as opposed to VPS4 and Rab8a.

**Figure 6.**
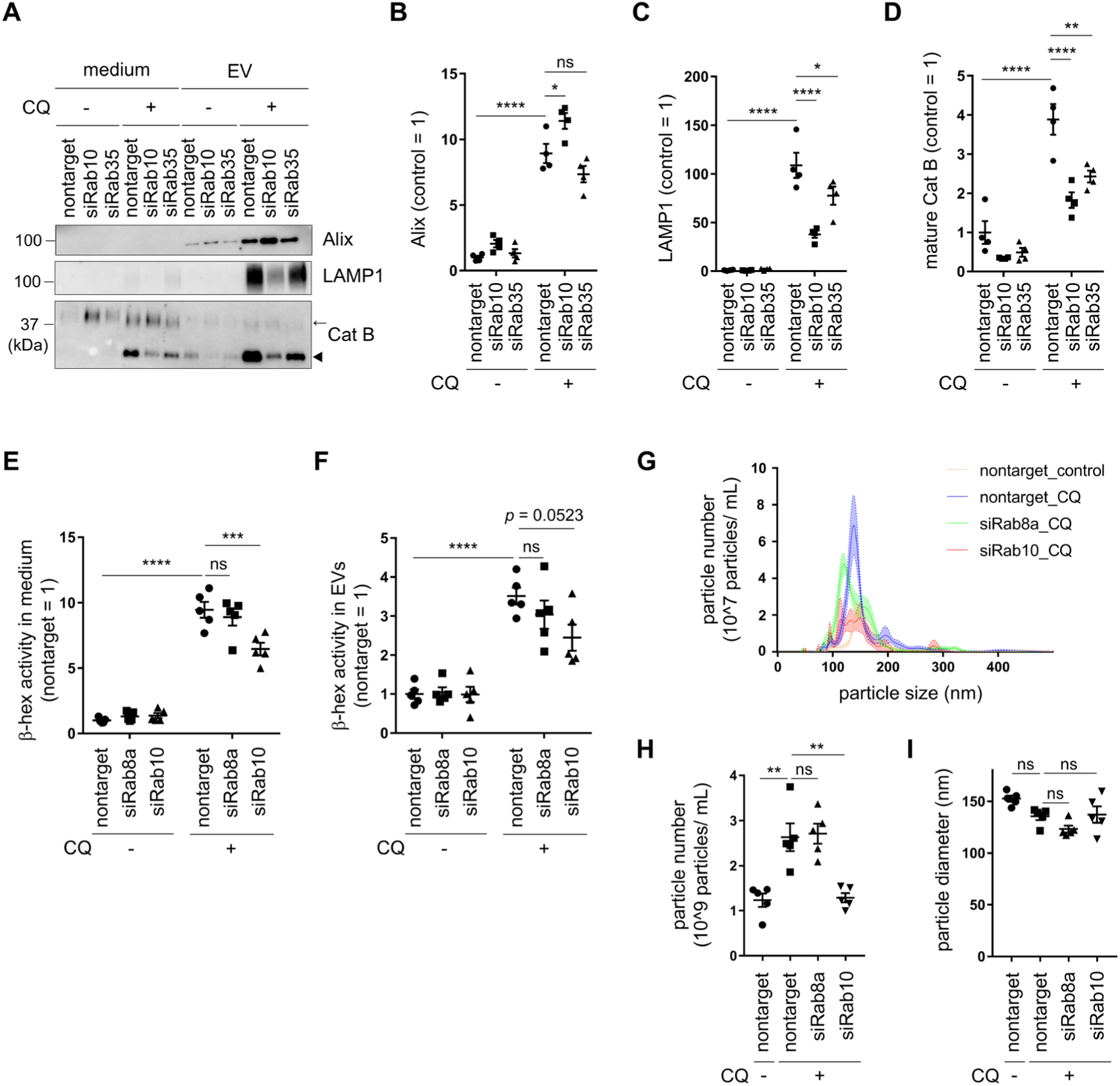
Rab10 and Rab35 regulate the secretion of LAMP1/cathepsin B-positive EVs under lysosomal stress. **(A)** Immunoblotting of medium and EV fractions derived from RAW264.7 cells treated with Rab10 or Rab35 siRNAs. Arrowhead: mature cathepsin B (Cat B), arrow: immature Cat B. **(B-D)** Quantification of Alix (B), LAMP1 (C) and mature Cat B (D) in EV fractions. n = 4, mean ± SEM, two-way ANOVA with Tukey’s test. * *p* < 0.05, ** *p* < 0.01, **** *p* < 0.0001. **(E, F)** β-Hex activity in medium (E) and EV (F) fractions derived from cells treated with Rab8a or Rab10 siRNAs. n = 5, mean ± SEM, one-way ANOVA with Tukey’s test. *** *p* < 0.001, **** *p* < 0.0001. **(G)** Size and number of the secreted EV particles measured by NTA. **(H, I)** Quantification of the number (H) and size (I) of particles detected in the culture media from cells under Rab8a or Rab10 knockdown conditions and treated with or without CQ, as shown in G. n = 5, mean ± SEM, one-way ANOVA with Tukey’s test. ** *p* < 0.01, **** *p* < 0.0001.

To further investigate whether such LAMP1/cathepsin B-positive EVs are of lysosomal origin, we assessed the β-hex activity in culture media and EV fractions of CQ-treated cells under Rab8a or Rab10 knockdown conditions. β-Hex activity in media decreased upon Rab10 knockdown, and its activity in EV fractions also decreased to a level close to significance (*p* = 0.0523), whereas Rab8a knockdown did not alter activity in either fraction **(Fig. 6E and 6F)**. This result suggests that β-hex is secreted along with LAMP1/cathepsin B-positive EVs via the Rab10 pathway.

We additionally examined the number and size of secreted EVs under Rab8a or Rab10 knockdown conditions by performing NTA. As a result, knockdown of Rab10, but not Rab8a, resulted in a significant decrease in the number of secreted EV particles **(Fig. 6G-H)**. The particle size remained unchanged on average **(Fig. 6I)**, whereas the knockdown of them altered the particle size distribution, resulting in the formation of multiple peaks **(Fig. 6G)**. The results support the notion that Rab8a and Rab10 regulate the secretion of some EV subtypes, and that Rab10 influences the total number of secreted EVs similarly to LRRK2.

### Syntaxin 2 and VAMP8 mediate the secretory process of multiple EV types downstream of LRRK2

To further clarify the mechanism leading to EV secretion downstream of LRRK2, we conducted targeted knockdown screening for genes that, when knocked down, cause a decrease in the secretion of lysosomal hydrolases upon CQ treatment. We first adopted the method of β-hex activity measurement in culture media, as this method is most simple and sensitive and thus suitable for screening. We selected 71 genes that are predicted to be involved in the intracellular transport or fusion of endolysosome-derived vesicles, or to interact with LRRK2 **(Table S1)**. Two positive control genes, *Lrrk2* and *Atg16l1*, were also included. Systematic knockdown of these genes in RAW264.7 cells identified 9 genes that affect the increase in β-hex activity in media upon CQ treatment **(Fig. 7A)**. Then, we repeated the experiments for 9 genes as a second screening, and found that knockdown of *Stx2*, encoding syntaxin 2, was the most effective in suppressing the secretion of β-hex **(Fig. 7B)**.

**Figure 7.**
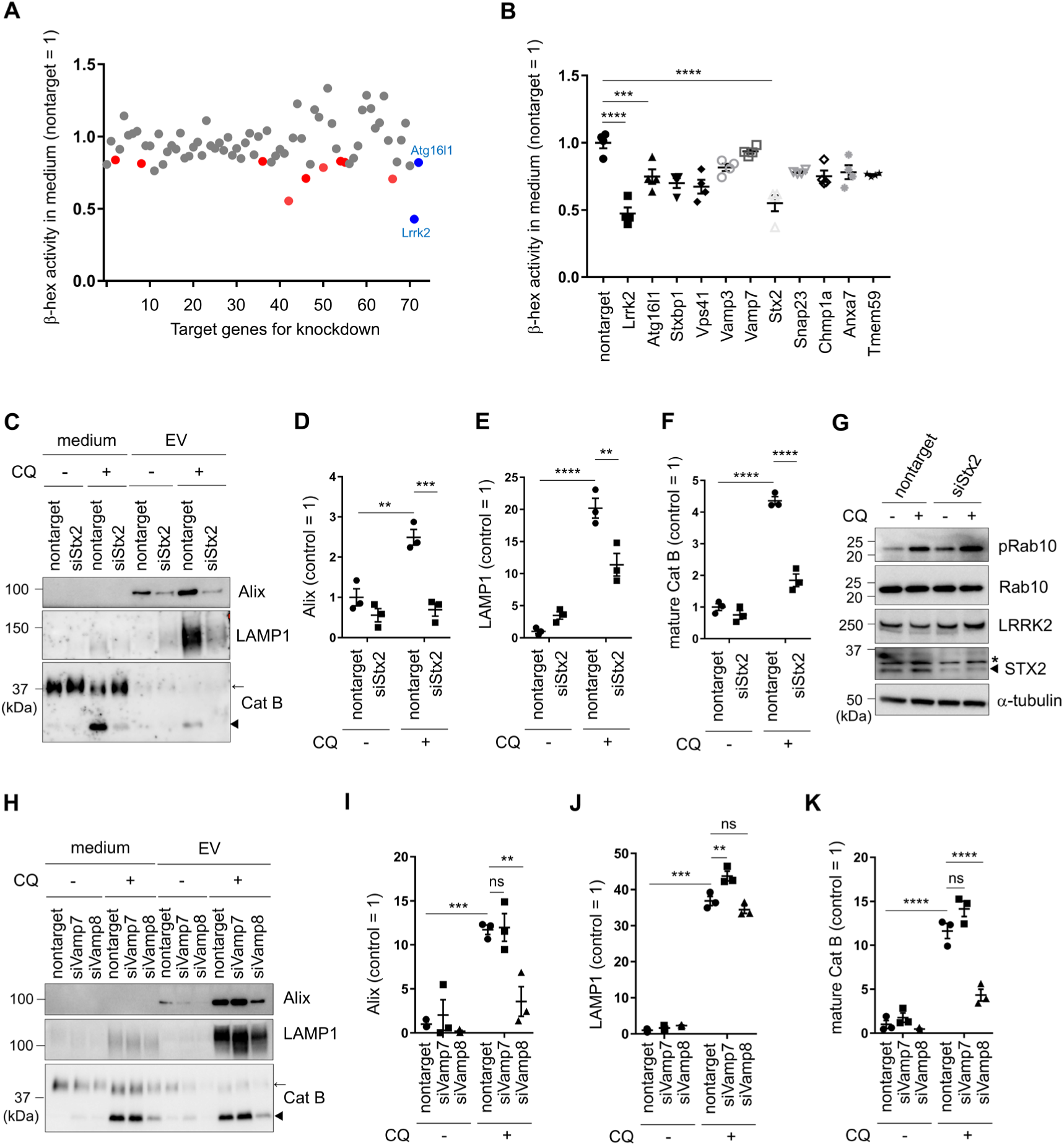
Syntaxin 2 and VAMP8 mediate the secretory process of multiple EV types downstream of LRRK2. **(A)** Knockdown screening for genes that suppress β-hexosaminidase (β-hex) activity in medium of RAW264.7 cells treated with CQ. The gene numbers on the horizontal axis correspond to the numbers in Table S1. Red: genes selected for 2nd screening. **(B)** β-Hex activity in medium of CQ-treated cells under knockdown of selected genes. n = 4, mean ± SEM, one-way ANOVA with Tukey’s test. *** *p* < 0.001, **** *p* < 0.0001. **(C)** Immunoblotting of medium and EV fractions derived from RAW264.7 cells treated with nontarget or *Stx2* siRNAs. Arrowhead: mature cathepsin B (Cat B), arrow: immature Cat B. **(D-F)** Quantification of Alix (D), LAMP1 (E) and mature Cat B (F) in EV fractions. n = 3, mean ± SEM, two-way ANOVA with Tukey’s test. ** *p* < 0.01, *** *p* < 0.001, **** *p* < 0.0001. **(G)** Immunoblot analysis of LRRK2 activity in cells treated with CQ under *Stx2* knockdown conditions. *: non-specific band. **(H)** Immunoblotting of medium and EV fractions derived from RAW264.7 cells treated with Vamp7 or Vamp8 siRNAs. Arrowhead: mature cathepsin B (Cat B), arrow: immature Cat B. **(I-K)** Quantification of Alix (I), LAMP1 (J) and mature Cat B (K) in EV fractions. n = 3, mean ± SEM, two-way ANOVA with Tukey’s test. ** *p* < 0.01, *** *p* < 0.001, **** *p* < 0.0001.

Syntaxin 2 is a SNARE protein on target membranes (t-SNARE) and is thought to regulate the final process of EV secretion by promoting membrane fusion with the plasma membrane. We thus examined the effect of *Stx2* knockdown on the secretion of each type of EVs. Knockdown of *Stx2* caused a decrease in the secretion of both Alix- positive EVs and LAMP1/cathepsin B-positive EVs upon CQ treatment, suggesting the involvement of syntaxin 2 in the secretion of EVs of both endosomal and lysosomal origins **(Fig. 7C-F)**. The levels of LRRK2 and phospho-Rab10 were not decreased by *Stx2* knockdown, indicating that syntaxin 2 does not regulate LRRK2 from upstream **(Fig. 7G)**. To further identify vesicle-associated SNARE (v-SNARE) that acts together with syntaxin 2, we focused on the candidate members of vesicle-associated membrane proteins (VAMPs), namely VAMP7 and VAMP8. *Vamp7* was included as a target gene in the above knockdown screening based on β-hex activity and had advanced up to the second screening, whereas *Vamp8* was not included. We examined the involvement of these proteins in the secretion of EVs under CQ treatment and found that knockdown of *Vamp8* significantly reduced the levels of Alix and mature cathepsin B in EV fractions **(Fig. 7H-K)**. Since VAMP8 has been reported as a partner of syntaxin 2 in the SNARE complex^53,54^, this result suggests that the VAMP8-syntaxin 2 interaction likely mediates the fusion of EVs with the plasma membranes upon secretion. Taking all results together, it has become clear that lysosomal stress induces the secretion of at least two types of EVs via the newly identified LRRK2-Rab-VAMP8-syntaxin 2 pathway, where Rab8a is involved in the secretion of Alix-positive EVs via ESCRT, while Rab10 and Rab35 are implicated in the secretion of lysosome-originated EVs **(Fig. 8)**.

**Figure 8.**
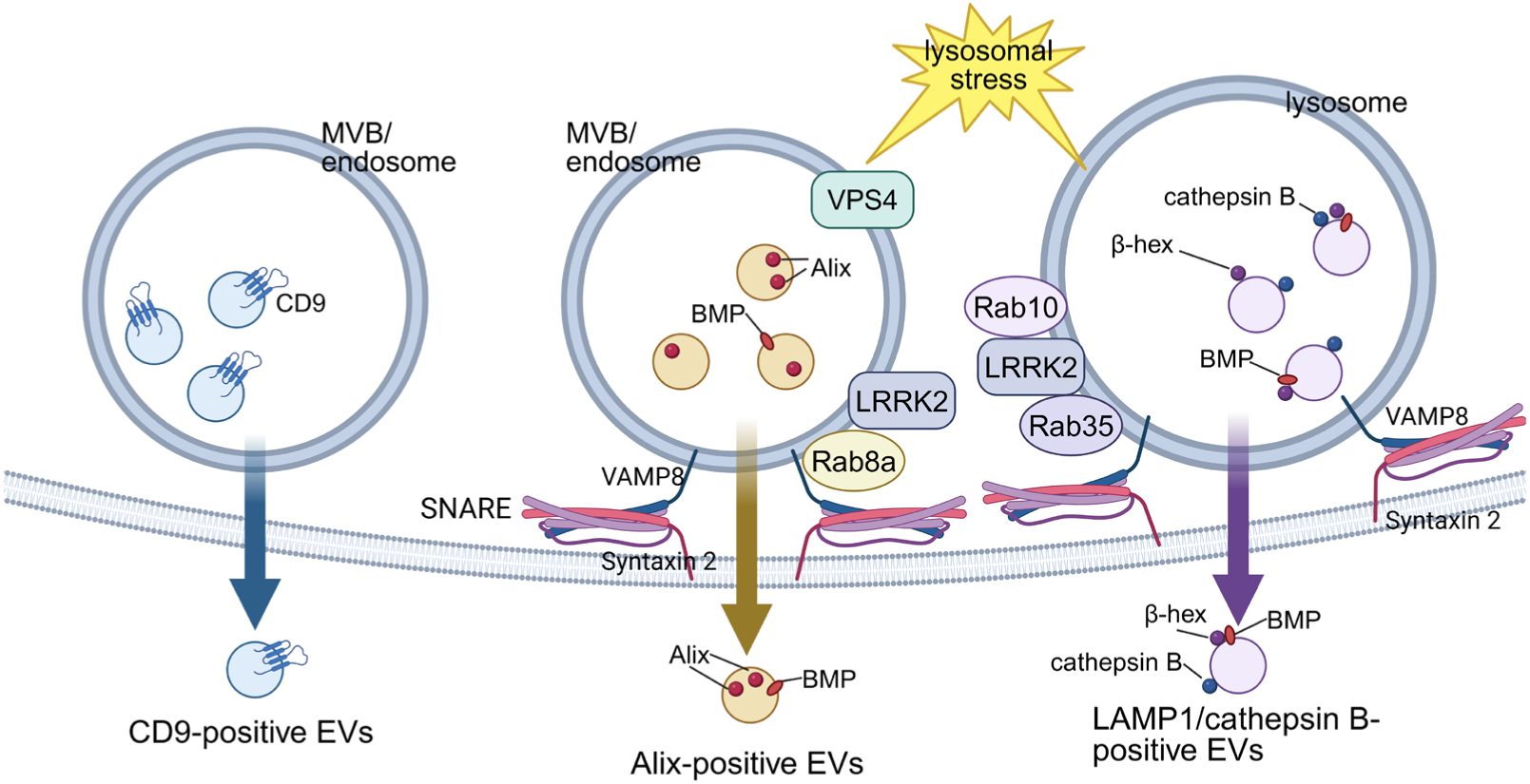
Schematic model of subtype-specific EV secretion by LRRK2 and substrate Rab GTPases. EVs are secreted via multiple pathways. CD9-positive EVs are secreted independently of lysosomal stress and LRRK2 (left). EVs secreted under lysosomal stress and LRRK2 regulation are BMP-positive, and their secretion is regulated by the VAMP8-syntaxin 2 SNARE complex. Furthermore, among LRRK2 substrates, Rab8a is involved in the secretion of Alix-positive EVs mediated by the ESCRT machinery (middle), while Rab10 and Rab35 are involved in the secretion of lysosome-derived EVs containing lysosomal hydrolases and LAMP1 (right). Secreted lysosomal hydrolases are primarily attached outside the EVs.

## Discussion

In this study, we revealed that the lysosomal stress elicits LRRK2-mediated secretion of EVs that exhibit unexpected diversity. EVs we detected in this study included at least three types: CD9-positive EVs, Alix-positive EVs, and LAMP1/cathepsin B-positive EVs, with the secretion of the latter two being both lysosomal stress-dependent and LRRK2-dependent. The observation that EVs had a diameter of 100-150 nm, along with detailed analyses on secretory pathways, supported the notion that the secreted EVs correspond to small EVs rather than microvesicles or secretory lysosomes. Also, secretion was enhanced in cells expressing PD-associated G2019S mutant LRRK2, and the endolysosomal lipid di-22:6-BMP, a urinary indicator elevated in LRRK2-associated and idiopathic PD, was also found to be secreted via this secretory pathway. These results raise the existence of a novel stress-responsive secretory mechanism that could also be involved in PD.

CQ and monensin used in this study are well-known lysosomal stressors, and it was reported over 40 years ago that these reagents induce the secretion of lysosomal enzymes from macrophages^55^. However, the mechanism behind this phenomenon has remained surprisingly undetermined. One explanation for this secretory phenomenon was that treatment with these reagents inhibited lysosomal function to degrade their contents, thereby expelling those contents outside the cell. Alternatively, as these reagents also upregulate autophagy markers, it has been considered that autophagosomes become unable to fuse with the impaired lysosomes, leading to the expulsion of their contents. Such secretion involving autophagy-related mechanisms is known to occur under certain conditions, a process also referred to as secretory autophagy during lysosome inhibition (SALI)^56,57^. However, our present study found no evidence of secretory autophagy, and electron microscopic observation of CQ-treated cells revealed very few double-membrane autophagosomes. Also, autophagy initiation complex components implicated in SALI (*e.g.,* FIP200) were not involved in this secretion process^34^. A similar secretory mechanism called LC3-dependent EV loading and secretion (LDELS) has also been reported^58,59^; however, we have confirmed that two representative cargos secreted by LDELS, SAFB and HNRNPK, were not detected in our experiments (unpublished observation). On the other hand, CQ and monensin are now known to induce CASM (conjugation of ATG8 to single membranes), not autophagy, and it is also clear that CASM activates LRRK2^34,60^ which then leads to the exocytic release of cathepsins^29,34^. Collectively, the actual mechanism of lysosomal enzyme and di-22:6-BMP secretion is thought to involve a pathway from lysosomal stress through CASM, LRRK2, its substrate Rab, and VAMP8-syntaxin 2, leading to EV release.

This study also revealed the diversity of EVs secreted under LRRK2 control. Specifically, such EVs constituted a subset of the total while including multiple types, and they were separately regulated by LRRK2 substrate Rabs. Although lysosomal stress may have further accentuated the diversity of EVs, such diversity has been noted in various studies to date^61^, and particular care is required when analyzing them collectively. EVs include ESCRT-dependent and ESCRT-independent types based on their biogenesis process, and our findings indicate that LRRK2 regulates ESCRT-dependent EVs only. This may be due to the roles of LRRK2-regulated Rabs, especially Rab8a, in the secretion of exosomes^51^ or their functional interaction with ESCRT or Alix^62^. Another finding that Rab10, not Rab8a, is involved in cathepsin secretion upon CQ treatment aligns with our previous report^29^, and now we additionally found that the cathepsin is secreted with EVs by attaching to their surface and that Rab35 is also involved in their secretion. The differences in the roles of Rab8a and Rab10/Rab35, as well as the coordinated function of Rab10 and Rab35, remain poorly understood and require further analyses. Our data also showed that secretion of LAMP1 and Cat B increased when the VPS4 pathway was inhibited, while secretion of Alix increased when the Rab10 pathway was inhibited. This result may indicate the existence of some compensatory relationship between these two pathways.

Understanding the LRRK2-mediated secretion mechanism reported here may also be helpful for elucidating the pathomechanism of PD. Another recent study has reported that neurons harboring LRRK2 G2019S mutation exhibit the secretion of EVs containing ATG8 proteins (*e.g.*, LC3, GABARAP) via secretory autophagy, with secretion increased by 15% compared to wild-type neurons^35^. On the other hand, the EV secretion phenotype we observed here differs in several aspects; it occurs primarily in macrophage lineage cells and rarely in neurons^33^, contains lysosomal hydrolases, and increases by 5-10-fold upon lysosomal stress loading. Our marker analysis and electron microscopy observations suggest that this lysosomal stress-induced secretion is not likely secretory autophagy, which is further supported by the fact that lysosomal stressors such as CQ induce CASM rather than autophagy^38,63^. Another key finding in the secretion phenotype reported here is that the endolysosomal phospholipid di-22:6-BMP is indeed secreted via this mechanism. Di-22:6-BMP is a biomarker of drug-induced phospholipidosis and lysosomal storage disorders, such as Niemann-Pick disease^64^. It is increased in the urine of PD patients with *LRRK2* gain-of function mutations (G2019S, R1441G/C), as well as retromer component *VPS35* D620N mutations, that amplify the LRRK2 response to endolysosomal stress^26–28^. Elevated urinary di-22:6-BMP is thought to be driven by increased exocytosis of BMP-enriched exosomes^36^. In addition, we have confirmed from our prior study that the secretion of lysosomal hydrolases observed here occurs in microglia as well^33^. Thus, together with the analysis of secretion from neurons, it would be crucial to conduct detailed analyses of changes in the secretion mechanisms within the brains or periphery of PD patients.

As a potential avenue for further identifying the links to PD pathomechanisms, investigation of the effects of secreted EVs at their destination sites would also be of interest. Among the secreted lysosomal hydrolases, a subset of cathepsins such as cathepsin B and S are known to retain hydrolase activity at near-neutral pH^65,66^ and thus have the potential to exert damaging effects upon reaching destinations. On the other hand, it has recently been reported that these cathepsins presumably attached on EVs can degrade extracellular α-synuclein, reducing the spreading of α-synuclein pathology^48^. From another perspective, we have reported that insoluble α-synuclein is released from microglia with EVs by this LRRK2-mediated mechanism upon CQ treatment^33^. How the EV secretion mechanism reported here contributes to the age-associated pathogenetic mechanisms including α-synuclein metabolism or propagation is an intriguing point, and it will be desired to verify the effect of such mechanisms *in vivo* in the future.

## Limitations of the study

First, this study was conducted under lysosomal stress conditions such as CQ treatment, and we need to further investigate the possible occurrence of similar secretion at low levels even under non-stressed conditions. Nonetheless, it is well known that signs of lysosomal stress, such as lysosomal enlargement and lipofuscin accumulation, become apparent in aged tissues^67,68^, and stressors like CQ may experimentally enhance such physiological changes associated with aging. Second, this study analyzed EV fractions recovered after filtration and ultracentrifugation of the culture medium, while other recovery methods such as affinity purification were also possible; we chose the ultracentrifugation method because it is still the most widely used technique. Third, we did not perform a proteome-wide analysis of secreted proteins, although simple proteomic analysis may not enable the selective detection of cleaved proteins such as mature cathepsins. Finally, this study did not include *in vivo* analyses of EV secretion. In particular, BMP is a urinary marker that reflects LRRK2 activity *in vivo*, and it is entirely plausible that the present mechanism is involved in the secretion of BMP into the urine. Given the connection to the aforementioned disease mechanisms, elucidating the role of EV secretion *in vivo* mediated by LRRK2 and lysosomal stress would be a key challenge for future research.

## Key resources table

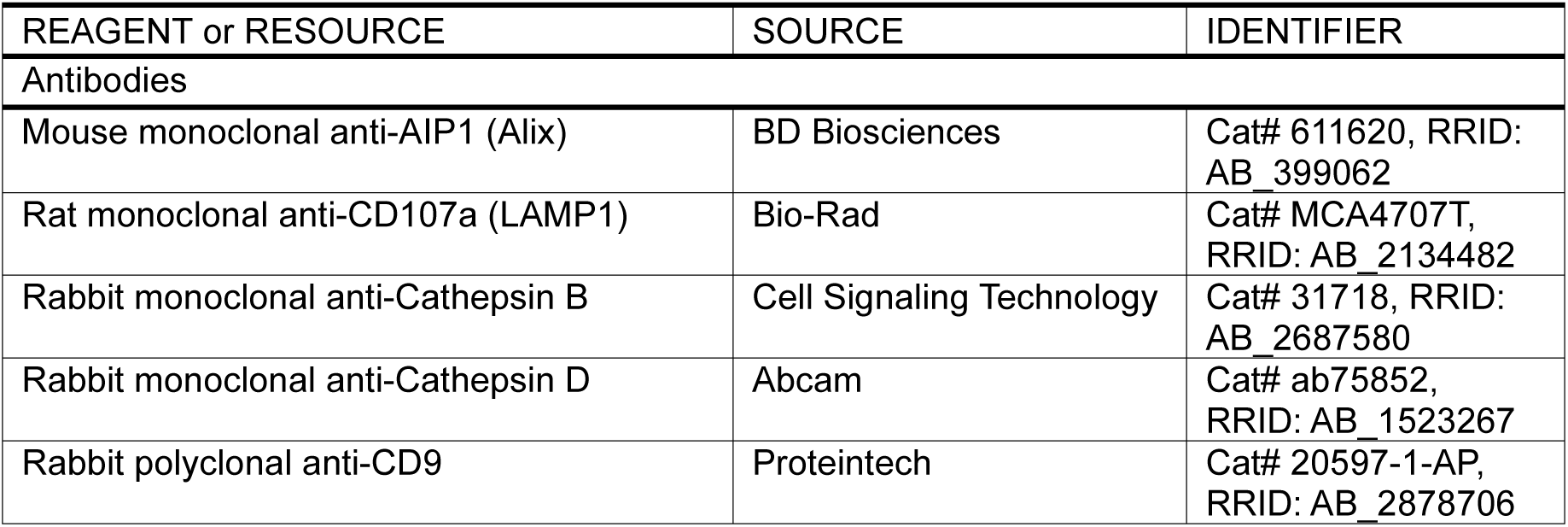
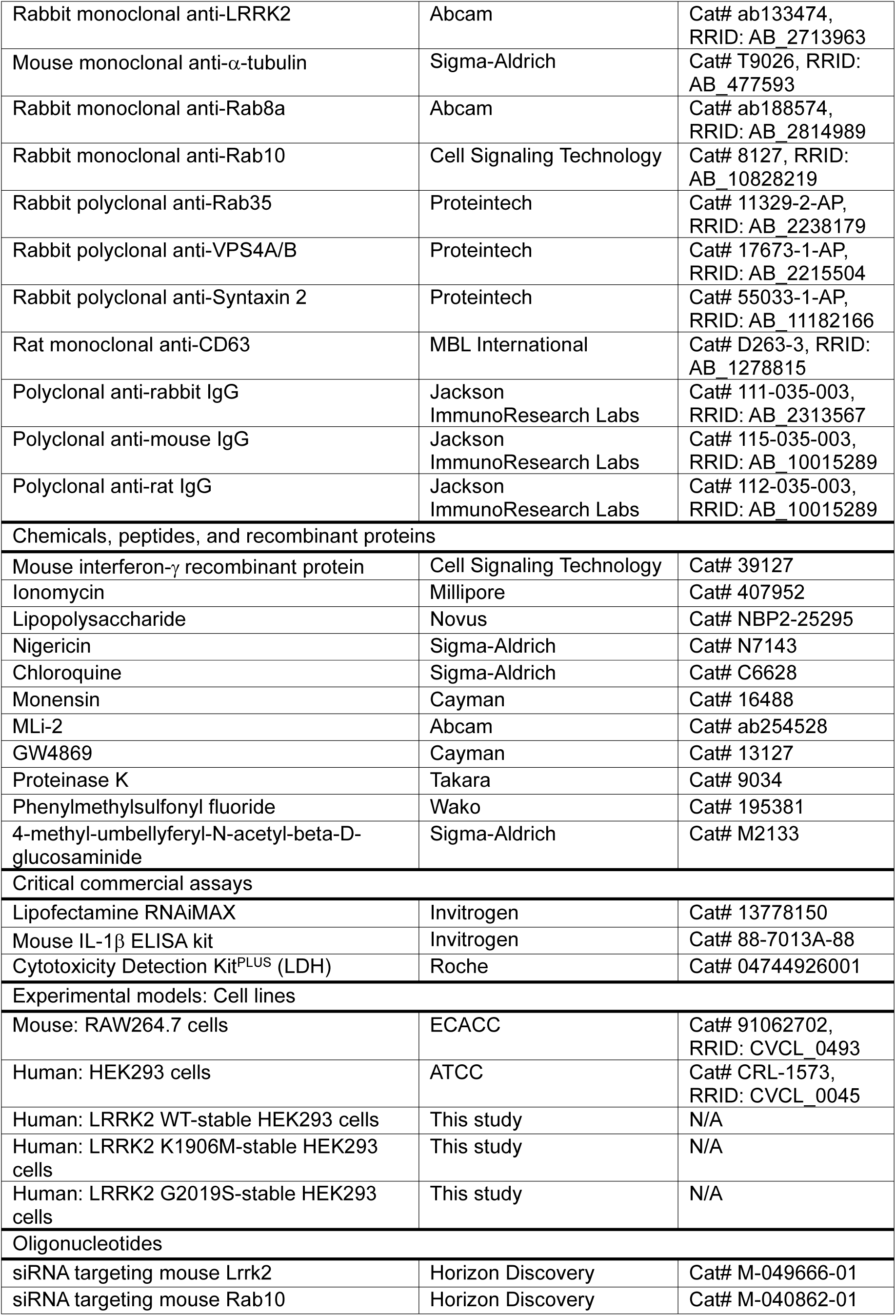
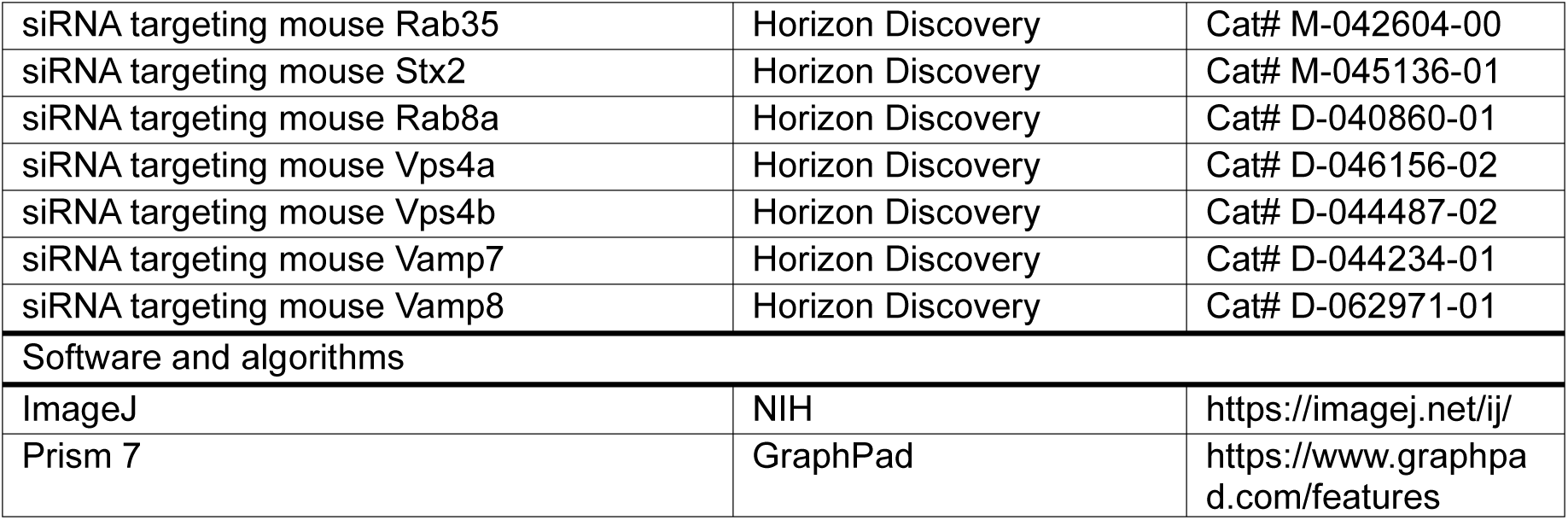

| REAGENT or RESOURCE | SOURCE | IDENTIFIER |
| --- | --- | --- |
| Antibodies |  |  |
| Mouse monoclonal anti-AIP1 (Alix) | BD Biosciences | Cat# 611620, RRID: AB_399062 |
| Rat monoclonal anti-CD107a (LAMP1) | Bio-Rad | Cat# MCA4707T, RRID: AB_2134482 |
| Rabbit monoclonal anti-Cathepsin B | Cell Signaling Technology | Cat# 31718, RRID: AB_2687580 |
| Rabbit monoclonal anti-Cathepsin D | Abcam | Cat# ab75852, RRID: AB_1523267 |
| Rabbit polyclonal anti-CD9 | Proteintech | Cat# 20597-1-AP, RRID: AB_2878706 |
| Rabbit monoclonal anti-LRRK2 | Abcam | Cat# ab133474, RRID: AB_2713963 |
| Mouse monoclonal anti-α-tubulin | Sigma-Aldrich | Cat# T9026, RRID: AB_477593 |
| Rabbit monoclonal anti-Rab8a | Abcam | Cat# ab188574, RRID: AB_2814989 |
| Rabbit monoclonal anti-Rab10 | Cell Signaling Technology | Cat# 8127, RRID: AB_10828219 |
| Rabbit polyclonal anti-Rab35 | Proteintech | Cat# 11329-2-AP, RRID: AB_2238179 |
| Rabbit polyclonal anti-VPS4A/B | Proteintech | Cat# 17673-1-AP, RRID: AB_2215504 |
| Rabbit polyclonal anti-Syntaxin 2 | Proteintech | Cat# 55033-1-AP, RRID: AB_11182166 |
| Rat monoclonal anti-CD63 | MBL International | Cat# D263-3, RRID: AB_1278815 |
| Polyclonal anti-rabbit IgG | Jackson ImmunoResearch Labs | Cat# 111-035-003, RRID: AB_2313567 |
| Polyclonal anti-mouse IgG | Jackson ImmunoResearch Labs | Cat# 115-035-003, RRID: AB_10015289 |
| Polyclonal anti-rat IgG | Jackson ImmunoResearch Labs | Cat# 112-035-003, RRID: AB_10015289 |
| Chemicals, peptides, and recombinant proteins |  |  |
| Mouse interferon-γ recombinant protein | Cell Signaling Technology | Cat# 39127 |
| Ionomycin | Millipore | Cat# 407952 |
| Lipopolysaccharide | Novus | Cat# NBP2-25295 |
| Nigericin | Sigma-Aldrich | Cat# N7143 |
| Chloroquine | Sigma-Aldrich | Cat# C6628 |
| Monensin | Cayman | Cat# 16488 |
| MLi-2 | Abcam | Cat# ab254528 |
| GW4869 | Cayman | Cat# 13127 |
| Proteinase K | Takara | Cat# 9034 |
| Phenylmethylsulfonyl fluoride | Wako | Cat# 195381 |
| 4-methyl-umbelliferyl-N-acetyl-beta-D-glucosaminide | Sigma-Aldrich | Cat# M2133 |
| Critical commercial assays |  |  |
| Lipofectamine RNAiMAX | Invitrogen | Cat# 13778150 |
| Mouse IL-1β ELISA kit | Invitrogen | Cat# 88-7013A-88 |
| Cytotoxicity Detection Kit<sup>PLUS</sup> (LDH) | Roche | Cat# 04744926001 |
| Experimental models: Cell lines |  |  |
| Mouse: RAW264.7 cells | ECACC | Cat# 91062702, RRID: CVCL_0493 |
| Human: HEK293 cells | ATCC | Cat# CRL-1573, RRID: CVCL_0045 |
| Human: LRRK2 WT-stable HEK293 cells | This study | N/A |
| Human: LRRK2 K1906M-stable HEK293 cells | This study | N/A |
| Human: LRRK2 G2019S-stable HEK293 cells | This study | N/A |
| Oligonucleotides |  |  |
| siRNA targeting mouse Lrrk2 | Horizon Discovery | Cat# M-049666-01 |
| siRNA targeting mouse Rab10 | Horizon Discovery | Cat# M-040862-01 |
| siRNA targeting mouse Rab35 | Horizon Discovery | Cat# M-042604-00 |
| siRNA targeting mouse Stx2 | Horizon Discovery | Cat# M-045136-01 |
| siRNA targeting mouse Rab8a | Horizon Discovery | Cat# D-040860-01 |
| siRNA targeting mouse Vps4a | Horizon Discovery | Cat# D-046156-02 |
| siRNA targeting mouse Vps4b | Horizon Discovery | Cat# D-044487-02 |
| siRNA targeting mouse Vamp7 | Horizon Discovery | Cat# D-044234-01 |
| siRNA targeting mouse Vamp8 | Horizon Discovery | Cat# D-062971-01 |
| Software and algorithms |  |  |
| ImageJ | NIH | https://imagej.net/ij/ |
| Prism 7 | GraphPad | https://www.graphpad.com/features |

## Experimental model and study participant details

### Cell lines

All cell lines used in this study are listed in this paper’s Key Resources Table. RAW264.7 cells (ECACC), HEK293 cells (ATCC) and NIH-3T3 cells were cultured in DMEM supplemented with 10% FBS and 100 U/mL penicillin and 100 mg/mL streptomycin at 37℃ in a 5% CO_2_ atmosphere. RAW264.7 cells were cultured on culture dish for suspended cells. RAW264.7 cells were activated by IFN-γ treatment (15 ng/mL) for 48 h before each assay. HEK293 cells stably expressing LRRK2 WT or mutants (G2019S, K1906M) were generated by transfecting cells with a plasmid encoding 3×FLAG-tagged full-length human LRRK2^18^ followed by selection under treatment with G-418 (Roche). Monoclonal stable cells were obtained after culturing in 96-well plates by the limiting dilution method.

## Method details

### Transfection and drug treatment

All siRNAs and chemicals used in this study are listed in Key Resources Table. Transfection of siRNA was performed using Lipofectamine RNAiMAX (Thermo Fisher Scientific) according to the manufacturer’s protocols. Cells were analyzed 48 h after siRNA transfection. The following reagents were added to cells by medium replacement at final concentrations as indicated: chloroquine (50-100 μM, 3 h), monensin (90 μM, 3 h), MLi-2 (100 nM, 3 h), GW4869 (100 μM, 3 h), ionomycin (2 μM, 3 h), nigericin (20 μM, 1 h).

### Measurement of β-hexosaminidase activity

The protocol was arranged from the method of β-hexosaminidase activity measurement in MEF cells^69^. RAW264.7 cells were seeded on 6-well plates at 25% confluency and then treated with IFN-γ as described above. 48 h later, cells were treated with or without 2 μM ionomycin (for analysis of lysosomal exocytosis), 100 μM CQ and 100 nM MLi-2 for 3 h. HEK293 cells were treated with 50 μM CQ for 3 h, and NIH-3T3 cells were treated with 50 or 100 μM CQ and 100 nM MLi-2 for 3 h. For analysis of EV fraction, EVs were collected from 1 mL of culture medium by ultracentrifugation by the following method and suspended in 80 μL PBS. 75 μL of culture medium or EV suspension were mixed with 25 μL of substrate buffer (4 mM 4-methyl-umbellyferyl-N-acetyl-beta-D-glucosaminide in 20 mM Na citrate-phosphate buffer, pH 4.0 (0.2 M Na_2_HPO_4_, 0.1 M citric acid)) in 96-well black plates, and the plates were incubated for 1 h at RT preventing from light. Then, 25 μL of stop solution (2 M Na_2_CO_3_, 1.1 M glycine) was added, and the fluorescence (Ex/Em = 365 nm/450 nm) was measured by using a plate reader (SpectraMax M2, WAKENYAKU).

### IL-1β ELISA

RAW264.7 cells were pre-treated with IFN-γ for 48 h and then with 500 ng/mL lipopolysaccharide (LPS) for 24 h. Cells were treated with or without 20 μM nigericin and 100 nM MLi-2 for 1 h, and culture medium was collected to perform ELISA using a mouse IL-1β ELISA kit (Invitrogen) according to the manufacturer’s protocols. Absorbance signals of each well at 450 nm and 570 nm were read by using a plate reader (SpectraMax M2, WAKENYAKU).

### Isolation of extracellular vesicles

RAW264.7 cells were cultured on 10 cm dishes under the conditions described above. The media were replaced with 5 mL of DMEM containing 1% FBS, 100 U/mL penicillin and 100 mg/mL streptomycin (for immunoblotting and β-hexosaminidase measurement) or those without FBS (for nanoparticle tracking analysis and BMP measurement) in the presence of the indicated reagents for 3 h. Then, the culture media were collected and centrifuged at 300 × *g* for 10 min at 4℃, and the supernatants were collected as “medium” fraction. This fraction was further centrifuged at 2,000 × *g* for 10 min at 4℃, and the supernatants were filtrated by using 0.22 μm filter (Millipore) to remove large vesicles. 4 mL of filtrated samples were ultracentrifuged at 210,000 × *g* for 70 min at 4℃ using TLA-110 rotator (Beckman-Coulter). The supernatants were collected as “sup” fraction. The pellets were rinsed with 2 mL of PBS and ultracentrifuged again in the same condition above. The supernatants were removed, and the pellets were suspended in 60 μL of PBS to obtain “EV” fractions.

### SDS-PAGE and immunoblot analysis

Detailed protocol of sample preparation for SDS-PAGE was described in our previous paper^70^. SDS-PAGE was performed by using 5-20% gradient gel (SuperSep Ace, 5-20%, FUJIFILM Wako). Transfer was performed by using iBlot3 Transfer system (Thermo Fisher). The transferred PVDF membranes were first blocked with 5% skim milk/0.1% Tween 20 in TBS and then incubated with the primary antibody diluted in Immunoenhancer (Wako) for 2 h at room temperature or overnight at 4℃. The membranes were washed with 0.1% Tween 20 in TBS and then incubated with the horseradish peroxidase (HRP)-conjugated secondary antibody for 45 min at room temperature. Chemiluminescent signals of the bands were excited by ImmunoStar Zeta reagent (Fujifilm) and were detected by LAS4000 (Fujifilm). Densitometric analysis of the bands was performed using ImageJ software (NIH).

### Negative staining of EVs

RAW264.7 cells cultured on 12-well plates were treated with 15 ng/mL IFN-γ for 36 h for priming. EVs were isolated from media by ultracentrifugation (100,000 × *g*, 70 min)^71^ and were resuspended in PBS. Samples were applied to Collodion film on the TEM grids (Nisshin-EM) and stained with 1% uranyl acetate for 3 min. After washing with ultrapure water, samples were observed with electron microscopy (JEOL, 1200EXⅡ).

### Nanoparticle tracking analysis

EVs collected by ultracentrifugation were suspended in 100 μL PBS. Particles were analyzed and calculated by NanoSight (Quantum Design), which was performed at Fujifilm Wako bio solutions.

### Proteinase K digestion

The protocol was arranged from Meneses-Salas et al^36^. The media collected from cell culture dishes were filtrated by using 0.22 μm filter (Millipore) and then incubated with Proteinase K (1/5000 (v/v)) (Takara) for 15 min on ice. Then, 1 μM phenylmethylsulfonyl fluoride (PMSF, Wako) was added to stop digestion, and EVs were isolated by ultracentrifugation as described above.

### Electron microscopic analysis of ILVs

RAW264.7 cells were cultured on cover glasses for 2 days in the presence of IFN-γ as described above. Cells were fixed with the solution containing 2% PFA, 2.5% Glutaraldehyde and 0.1M phosphate buffer (PB) for 1 h at RT and then washed with 0.1M PB. Cells were incubated with 0.1M Cacodylate-HCl (pH 7.4) and then post-fixed in reduced osmium (1% OsO_4_, 1.5% potassium ferrocyanide in 0.1M cacodylate buffer) for 1 h on ice. After washing with water, cells were rinsed with 70% EtOH and then washed for 10 min with 80%, 90%, 95% and 100% EtOH (twice) for dehydration. Cells were incubated with QY-1 (Nisshin-EM) for 10 min twice and then rotated with Durcupan (Sigma-Aldrich) plus QY-1 (1:1) overnight at room temperature. Cells were rotated with 100% Durcupan for 2 h twice and cover glasses were embedded in fresh Durcupan and incubated 48 h at 60℃. Cover glasses were removed from resin blocks using liquid nitrogen. Ultrathin sections (50-70 nm) were prepared using an ultramicrotome (Leica, REICHERT NISSEI, Ultracut S) and picked up onto formvar-coated grids (Okenshoji). The sections were stained with 4% uranyl acetate for 5 min in the dark under humid conditions at room temperature. After washing, the sections were treated with Reynolds lead citrate in the presence of solid KOH for 2 min at room temperature, washed with water and air-dried. The sections were imaged using TEM (JEOL, 1200EXⅡ) operating at 80-90 kV and equipped with a CCD camera (JEOL, Veleta).

### Measurement of BMP levels in cell pellets and EVs

RAW264.7 cells cultured on 10 cm dishes were treated with the indicated reagents, and EVs were collected from culture media by ultracentrifugation as described above followed by suspension in 50 μL DPBS (Gibco). The cells were collected using cell scraper and centrifuged 300 × *g* for 5 min at 4℃ to obtain cell pellets. EVs and cell pellets were temporally frozen at -80℃. The quantitation of di-22:6 BMP in EVs and cell pellets was conducted at Nextcea, Inc. (Woburn, MA) as previously described^27,64^ using a Shimadzu Nexera XR UPLC system (Shimadzu Scientific Instruments, Japan) and a Sciex 7500 mass spectrometer (Sciex, USA). The di-22:6 BMP standard curve was prepared. Internal standard di-22:6 BMP-d5 was used.

### Statistics

Statistical analysis was performed using GraphPad Prism 7 (GraphPad software). We adopted one-way ANOVA or two-way ANOVA for comparisons among three or more groups, and unpaired two-tailed *t*-test for comparisons between two groups. Dunnett’s test and Tukey’s test were used for post-hoc test after one-way and two-way ANOVA. Standard error of the mean (SEM) was shown in all graphs. *P* values less than 0.05 were considered statistically significant. No exclusion criteria were applied to exclude samples from analysis.

## Resource availability

### Lead contact

Requests for further information and resources and reagents should be directed to and will be fulfilled by the lead contact, Tomoki Kuwahara.

### Materials availability

This study did not generate new unique reagents.

### Data and code availability

- All data reported in this paper will be shared by the lead contact upon request.
- This paper does not report original code.
- Any additional information required to reanalyze the data reported in this work paper is available from the lead contact upon request.

## Acknowledgements

The authors thank the Iwatsubo lab members for helpful suggestions and discussions. This work was supported by JSPS KAKENHI grant numbers 20H00525 (T. I.), 21J12881 (M. S.), 22H02949 (T. K.), 23K19397 (M. S.), 24K18616 (M. S.), 25K02458 (T. K.) and by Takeda Science Foundation research grant (T. K.).

## Author contributions

Conceptualization: M. S. and T. K.; Investigation: M. S., T. K., S. S., T. S., E. T.; Methodology: M. S., S. T. and F. H.; Resources: S. T., T. T. and F. H.; Data curation: M. S., T. K. and F. H.; Writing – original draft: M. S. and T. K.; Writing – review & editing: T. K., F. H. and T. I.; Supervision: T. K. and T. I.; Project administration: T. K.; Funding acquisition: M. S., T. K. and T. I. All authors read and approved of the final manuscript.

