## Supplemental Figures for "Subtype-specific secretion of extracellular vesicles by LRRK2 and Rab GTPases under lysosomal stress"

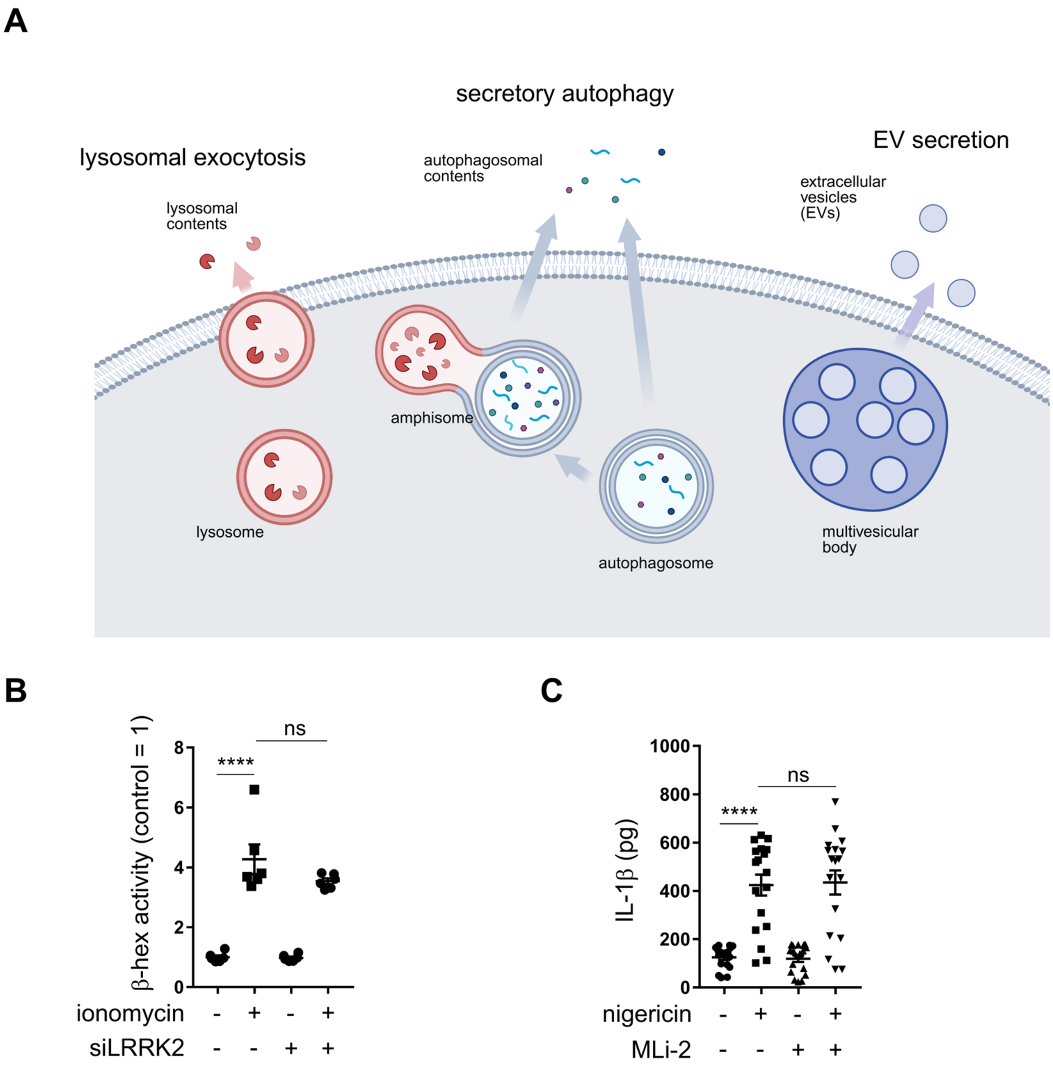


**Figure S1. Analysis of LRRK2 involvement in the known endolysosomal secretory pathways, related to Figure 1.**

**(A)** Three known endolysosomal secretory pathways that differ in membrane dynamics but are suggested to involve ATG8 conjugation system proteins acting upstream of LRRK2. Created in BioRender. Sakurai, M. (2026) https://BioRender.com/6u7hyvn **(B)** Assessment of lysosomal exocytosis by measuring β-hex activity in the medium of RAW264.7 cells pretreated with siLRRK2 and then treated with ionomycin. n = 6, mean ± SEM, one-way ANOVA with Tukey's test. **** $p <$ 0.0001. **(C)** Assessment of secretory autophagy by measuring IL-1b levels in the medium of RAW264.7 cells treated with or without nigericin and MLi-2. n = 19, mean ± SEM, one-way ANOVA with Tukey's test. **** $p < 0.0001$.

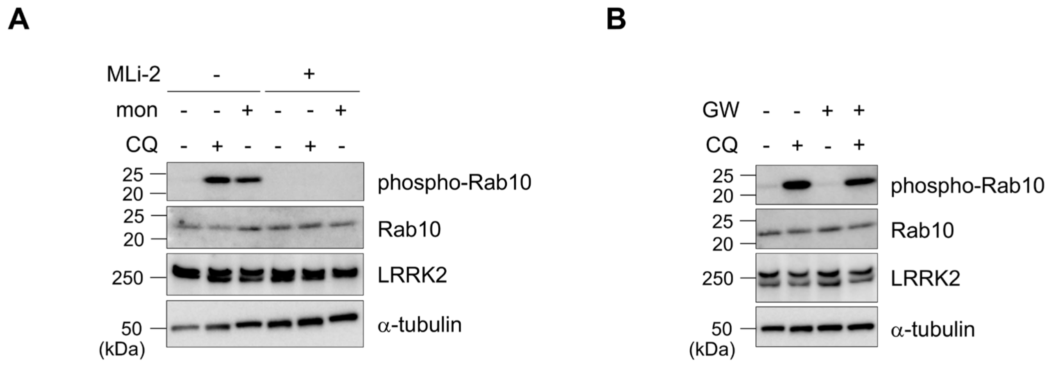


**Figure S2. LRRK2 activation upon CQ or monensin treatment and its independence from GW4869 treatment, related to Figures 1 and 2.**

**(A)** Immunoblot analysis of cell lysates derived from RAW264.7 cells treated with or without CQ, monensin and MLi-2. **(B)** Immunoblot analysis of cell lysates derived from RAW264.7 cells treated with or without CQ and GW4869.

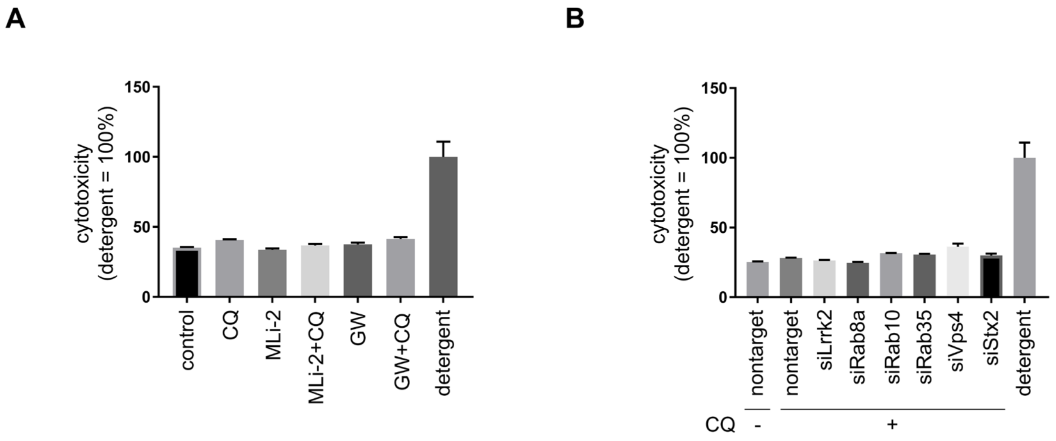


**Figure S3. Lack of cytotoxicity following drug treatment or knockdown, as assessed by LDH activity measurement, related to Figures 1, 2 and 5-7.**

**(A)** Cytotoxicity was assessed by measuring LDH activity in culture media of RAW264.7 cells treated with CQ, MLi-2 or GW4869. n = 3, mean ± SEM. **(B)** Cytotoxicity was assessed by measuring LDH activity in culture media of RAW264.7 cells treated with the indicated siRNAs as well as CQ. n = 3, mean ± SEM.

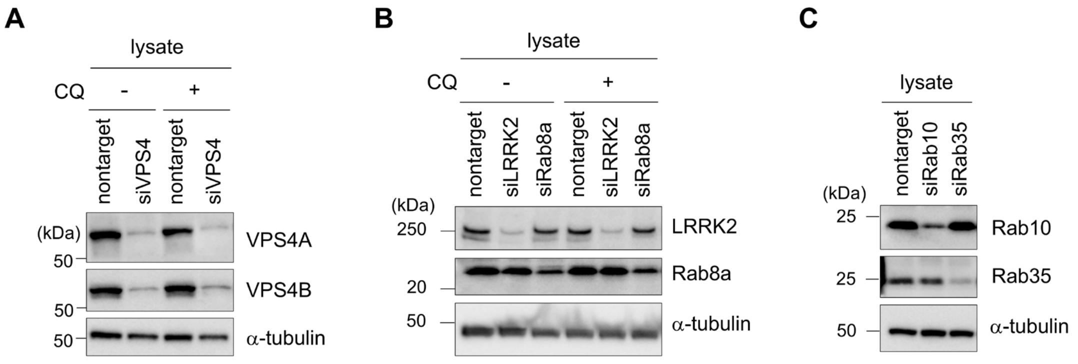


**Figure S4. Confirmation of the knockdown efficiencies, related to Figures 5 and 6.**

**(A-C)** Immunoblot analysis of protein knockdown efficiencies of VPS4A/B (A), LRRK2 (B), Rab8a (B), Rab10 (C) and Rab35 (C) in RAW264.7 cells treated with each siRNA as indicated.

**Table S1. List of genes targeted for knockdown screening, related to Figure 7.**

| No. | gene | protein |
|---|---|---|
| 1 | Acbd3 | acyl-CoA binding domain containing 3 |
| 2 | Alg2 | ALG2 alpha-1,3/1,6-mannosyltransferase |
| 3 | Anxa7 | annexin A7 |
| 4 | Arf6 | ARF GTPase 6 |
| 5 | Arpc2 | actin-related protein 2/3 complex subunit 2 |
| 6 | Arpc4 | actin-related protein 2/3 complex subunit 4 |
| 7 | Ccdc93 | CCC complex scaffolding subunit CCDC93 |
| 8 | Cdc42 | cell division cycle 42 |
| 9 | Chmp1a | charged multivesicular body protein 1A |
| 10 | Chmp3 | charged multivesicular body protein 3 |
| 11 | Chmp6 | charged multivesicular body protein 6 |
| 12 | Clint1 | clathrin interactor 1 |
| 13 | Coro1b | coronin 1B |
| 14 | Coro1c | coronin 1C |
| 15 | Coro7 | coronin 7 |
| 16 | Dnajc5 | DnaJ heat shock protein family (Hsp40) member C5 |
| 17 | Eea1 | early endosome antigen 1 |
| 18 | Ehd1 | EH domain containing 1 |
| 19 | Ehd4 | EH domain containing 4 |
| 20 | Fam129a | niban apoptosis regulator 1 |
| 21 | Fam129b | niban apoptosis regulator 2 |
| 22 | Fam134b | reticulophagy regulator 1 |
| 23 | Fam184b | family with sequence similarity 184 member B |
| 24 | Fam65c | RIPOR family member 3 |
| 25 | Gopc | golgi associated PDZ and coiled-coil motif containing |
| 26 | Gphn | gephyrin |
| 27 | Hgs | hepatocyte growth factor-regulated tyrosine kinase substrate |
| 28 | Mapk8ip3 | mitogen-activated protein kinase 8 interacting protein 3 |
| 29 | Mvb12a | multivesicular body subunit 12A |
| 30 | Myo1d | myosin ID |
| 31 | Nsf | N-ethylmaleimide sensitive factor, vesicle fusing ATPase |
| 32 | Pacsin2 | protein kinase C and casein kinase substrate in neurons 2 |
| 33 | Pdcd6ip | programmed cell death 6 interacting protein |
| 34 | Rac1 | Rac family small GTPase 1 |
| 35 | Scamp5 | secretory carrier membrane protein 5 |
| 36 | Sec22a | SEC22 homolog A, vesicle trafficking protein |
| 37 | Snap23 | synaptosome associated protein 23 |
| 38 | Snf8 | SNF8 subunit of ESCRT-II |
| 39 | Snx27 | sorting nexin 27 |
| 40 | Snx6 | sorting nexin 6 |
| 41 | Spag9 | sperm associated antigen 9 |
| 42 | Sqstm1 | sequestosome 1 |
| 43 | Stx2 | syntaxin 2 |
| 44 | Stx3 | syntaxin 3 |
| 45 | Stx4a | syntaxin 4 |
| 46 | Stx6 | syntaxin 6 |
| 47 | Stxbp1 | syntaxin binding protein 1 |
| 48 | Syt5 | synaptotagmin 5 |
| 49 | Syt7 | synaptotagmin 7 |

| | | |
|---|---|---|
| 50 | Tmem175 | transmembrane protein 175 |
| 51 | Tmem59 | transmembrane protein 59 |
| 52 | Tsg101 | tumor susceptibility 101 |
| 53 | Usp19 | ubiquitin specific peptidase 19 |
| 54 | Vamp2 | vesicle associated membrane protein 2 |
| 55 | Vamp3 | vesicle associated membrane protein 3 |
| 56 | Vamp7 | vesicle associated membrane protein 7 |
| 57 | Vps11 | VPS11 core subunit of CORVET and HOPS complexes |
| 58 | Vps16 | VPS16 core subunit of CORVET and HOPS complexes |
| 59 | Vps25 | vacuolar protein sorting 25 homolog |
| 60 | Vps26b | VPS26 retromer complex component B |
| 61 | Vps28 | VPS28 subunit of ESCRT-I |
| 62 | Vps29 | VPS29 retromer complex component |
| 63 | Vps35 | VPS35 retromer complex component |
| 64 | Vps36 | vacuolar protein sorting 36 homolog |
| 65 | Vps37a | VPS37A subunit of ESCRT-I |
| 66 | Vps39 | VPS39 subunit of HOPS complex |
| 67 | Vps41 | VPS41 subunit of HOPS complex |
| 68 | Vps51 | VPS51 subunit of GARP complex |
| 69 | Vti1b | vesicle transport through interaction with t-SNAREs 1B |
| 70 | Washc4 | WASH complex subunit 4 |
| 71 | Wdr91 | WD repeat domain 91 |
| **Positive controls** | | |
| 72 | Lrrk2 | leucine rich repeat kinase 2 |
| 73 | Atg16l1 | autophagy related 16 like 1 |
